# Aggregation in experimental studies with microparticles: Bacterial communities in the exposure system affect animal responses to the test particles

**DOI:** 10.1101/2025.09.13.676006

**Authors:** Sophia Reichelt, Rehab El-Shehawy, Elena Gorokhova

## Abstract

The role of microorganisms is frequently overlooked in effect studies with particulate materials, such as microplastics. In addition to the microbes naturally found in the environment, test animals can transfer their microbiome to the surrounding media and establish bacterial communities in the exposure vessels. The interactions between the animals and the bacterial communities during the exposure can influence the animal responses to experimental factors, such as particle abundance, aggregation, and other characteristics. However, the current designs in particle ecotoxicology often overlook these interactions.

In our 72-hour experiment, *Daphnia magna* were exposed to mixed kaolin clay and microplastics (<20-µm polystyrene fragments). We aimed to assess microbial communities derived from *Daphnia* microbiota, focusing on particle-associated biofilms and non-adherent cells and the effects of the total suspended solids (1-10 mg/l), microplastics contribution (0-10%), dissolved organic matter (agarose; 0 and 20 mg/l), and aggregate size/topology on these communities. Furthermore, we explored the impact of bacterial diversity and community composition on *Daphnia* mortality and body condition using individual protein content as a proxy.

We found a high similarity between bacterial communities and the *Daphnia* microbiome, indicating the microbiome as the source. Experimental factors had differential effects on the biofilms and non-adherent cells, with total suspended solids and agarose mainly influencing non-adherent cells at the family level (mostly upregulation) and microplastics affecting biofilms (both up- and downregulation). Aggregate size and topology were the key predictors of bacterial alpha diversity and the abundance of the affected families. Finally, the adverse effects on *Daphnia* were primarily driven by small aggregate size, agarose addition, and high biofilm diversity. These findings underscore the need to consider microbial components and their interactions with particles and species to comprehensively understand microplastic effects and develop ecologically relevant hazard assessment assays.

## 1. Introduction

The widespread presence of microplastics (MP) in various environmental compartments has sparked much debate regarding their impact on organisms, from microbes to whales. In the last decade, there has been a considerable increase in MP effect studies conducted on aquatic species. These studies employ a wide range of endpoints, including subcellular responses, changes in energy storage, feeding, growth and reproduction, as well as increased mortality and transgenerational effects (Doyle et al., 2022; Schür et al., 2020; Sun et al., 2021). However, the findings are frequently conflicting, with adverse effects, lack of effects, or even increased feeding and growth commonly observed in response to MP exposure (Bucci et al., 2019). It is widely acknowledged that conducting a comprehensive and ecologically relevant evaluation of MP impacts is a complex task, primarily due to methodological limitations. The existing exposure systems face difficulties in effectively addressing the diverse nature and physicochemical properties of test particles and accurately replicating environmental conditions within laboratory settings (Koelmans et al., 2022; Rochman et al., 2019). Additionally, the planktonic and benthic test organisms possess varying capacities to adapt to particulate matter in their environment, further complicating the assessment process.

The observable effects of MP are often similar to those of noncaloric natural particles, such as clay or cellulose (Ogonowski et al., 2018). Clogging feeding appendages, food dilution with adverse effects on energy reserves, translocation to the internal tissue in heterotrophs, adherence to cell walls and membranes, and thus disrupting light harvesting in autotrophs (Prata et al., 2019) are examples of such effects. Nevertheless, notable plastic-related effects exist, particularly concerning chemical exposure to leachates, including adverse impacts occurring at environmentally relevant concentrations. For instance, there have been documented cases of acute toxicities in Coho salmon caused by a transformation product of a tire antioxidant (Tian et al., 2021).

Microplastics constitute a small fraction of naturally occurring particles, such as detritus, sediments, and black carbon. Therefore, to assess thier environmental impacts accurately, it is essential to consider the entire particle load in the system, which varies between high-turbidity eutrophic and low-turbidity oligotrophic environments. Moreover, most particles, including MP, form aggregates in the water column and sediments, driving particle size distribution (PSD) in the environment; this process is influenced by abiotic (particle size, organic matter, salinity, water flow) and biotic (primary producers, biofilms, animal activity) factors. Aggregation can be affected by MP abundance and properties, altering their accessibility for organisms (Reichelt and Gorokhova, submitted; Motiei et al., 2021). In particular, smaller MP fragments have been found to have a higher potential for direct interactions with feeding appendages and guts of aquatic filtrators, such as many model species used in ecotoxicity testing. However, enhanced aggregation can promote particle sedimentation and change particle accessibility for consumers, thereby impacting the direct effects of MP on test organisms in experimental settings (Thornton Hampton et al., 2022).

Interactions between microorganisms play a key role in driving particle aggregation in the environment. Microorganisms are an essential component of aggregates, and microbial biofilms on particle surfaces have multiple functions. Firstly, they represent a biota that is exposed to plastic litter and potentially impacted by this exposure (Yang et al., 2020). Secondly, they influence the distribution and fate of plastics in the environment, including the formation of aggregates (Sooriyakumar et al., 2022). Thirdly, they can mediate MP effects via adding caloric value to otherwise noncaloric particles, which can increase the nutritional appeal of the aggregate colonised by the biofilm, potentially raising the likelihood of ingestion by consumers (Jeong et al., 2016).

On the one hand, aggregation is facilitated by microbes, but on the other hand, the micro-scale heterogeneity due to variations in the aggregate size and topology drives the diversity and lifestyle choices of the microorganisms inhabiting these environments (Stocker, 2012). As a result, some microorganisms remain in the water phase as non-adherent cells, while others settle and form biofilms (Cai, 2020), surface-attached communities that offer protection from environmental stressors, facilitate cell-to-cell communication and the exchange of nutrients (Flemming et al., 2016). Moreover, most bacterial taxa switch continuously from a planktonic lifestyle to living in a biofilm. However, aggregate composition, including the presence of MP, can affect this dynamic equilibrium, with downstream effects on the microbial communities, aggregation dynamics, and multicellular organisms interacting with these aggregates in the water column.

Biofilm formation on plastic litter is mostly studied in field settings, and we know much less about its importance in laboratory experiments designed to test the effects of solid particles, such as MP, on plants and animals. In such experiments, the primary focus is often on the test organisms and their responses, whereas the microbial component and particle aggregation in the exposure system are not acknowledged. It has been shown, however, that microbial communities associated with test organisms quickly inoculate the system and become established in the experimental media, colonising test particles and affecting their aggregation (ref). Therefore, even when good laboratory practice is in place, and sterile media and utensils are used in the experiment, the biofilms and non-adherent bacteria are present, and their effects on the test outcome (i.e., the response of the test organism) must be understood.

The study objectives were to (1) assess the microbial community originating from the microbiota of the test organism *Daphnia magna* in the exposure system and present during the exposure as a biofilm on the particle aggregates and non-adherent cells in the media, (2) understand how experimental conditions, including test particles, aggregate PSD, and natural organic matter drive microbial diversity, and (3) evaluate whether the microbial component affects how the test species *Daphnia magna* is responding to the exposure.

## 2. Material and Methods

### 2.1 Test particles

We used polystyrene (PS; Goodfellow GmbH, density 1.05 g cm^-3^ [pristine polymer]) artificially aged with UV-light and fragmented to < 20 µm as a test MP and kaolin clay (Sigma-Aldrich, K7375; density 2.6 g cm^-3^; size < 40 µm) as a reference material (see Supplementary Information, Table S1, for details). Detailed descriptions of the PS preparation and the particle mixture preparation are presented by Gewert et al. (2018) and Gerdes et al., (2019), respectively. The stock suspensions were prepared using the *Daphnia* culture media M7 (Samel et al., 1999), also used in all experimental mixtures.

We measured PSD in the particle stocks and all test mixtures using Spectrex Laser Particle Counter (LPC; Spectrex PC-2000, Redwood, City, USA). The target size and count number linear ranges were 1-100 µm and 50-1000 particles/cc, respectively. The GRADISTAT software 9.1 (Blott and Pye, 2001) was used to derive the PSD parameters following the computational method by Folk and Ward (Folk and Ward, 1957) to obtain mean particle size, median particle size (D_50_), D_90_ (particle size below which 90% of particles are found), D_10_ (particle size below which 10% of particles are found), skewness and kurtosis of each PSD.

### 2.1 Test species

*Daphnia magna* (Clone 5; The Federal Environment Agency, Berlin, Germany) was used as a test organism. The daphnids were cultivated in M7 media at 20°C and fed three times a week with green algae *Pseudokirchneriella subcapitata* and *Scenedesmus subspicatus.* One-week-old *Daphnia* (2-2.5 mm; Instar III-IV) were isolated from the culture and starved overnight before the experiment. No reproduction occurred during the exposure.

### 2.2 Experimental set-up

#### 2.2.1 Aggregate generation

In preparation for the exposure experiment, we generated MP-kaolin aggregates using particle mixtures (Table S1 Supplementary Information, Table 1); see Reichelt and Gorokhova (submitted) for details. In brief, the suspended solids (SS; 0-10 mg/l) were mainly comprised of kaolin, with the polystyrene (MP%) contributing 0-10% by mass and 0-17% by the particle number (Table S1, Supplementary Information). These MP concentrations are representative of hot-spot areas, such as lakes and wastewater effluents; for example, 0.1-10 mg/L and 10^3^-10^7^ particles/m^3^ (Di and Wang, 2018; Lenz et al., 2016; Sun et al., 2019; Wang et al., 2018).

**Table 1.**
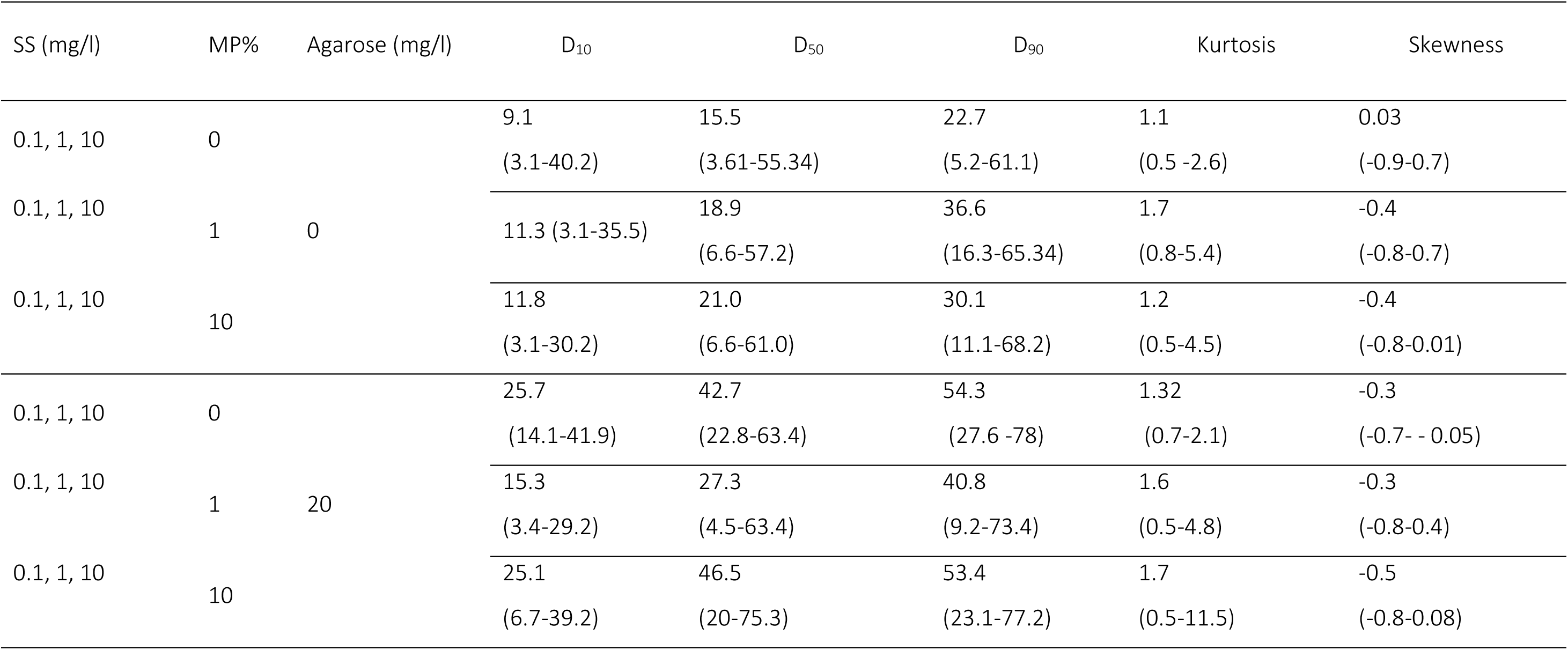
Overview of the experimental treatments. Total suspended solids (SS concentration; 0.1-10 mg/l) contained 0-10% microplastics (MP%), and agarose was added to half of the experimental units. The PSD in each experimental unit was assessed as D_10_, D_50_, D_90_, kurtosis and skewness; the average values and the range for each treatment (in parentheses) are shown.

Nine test mixtures representing different SS × MP% combinations and a particle-free media control were prepared using M7 media (Table S2). The same SS × MP% combinations were repeated, adding 20 mg/l of agarose as dissolved organic matter (DOM); this concentration is representative of high DOM levels in surface waters and sufficient to facilitate particle coating to mimic eco-corona.

#### 2.2.2 Experimental procedure

The incubation was carried out in scintillation vials, sealed airtight and mounted on a plankton wheel., The particle mixtures were first incubated for 48 h to allow for aggregate formation, and then daphnids were introduced into the vials (5 daphnids per vial; Figure 1). The vials were sealed and incubated on the plankton wheel for another 72 h following the experimental protocol of Gerdes and co-workers (Gerdes et al., 2019).

**Figure 1.**
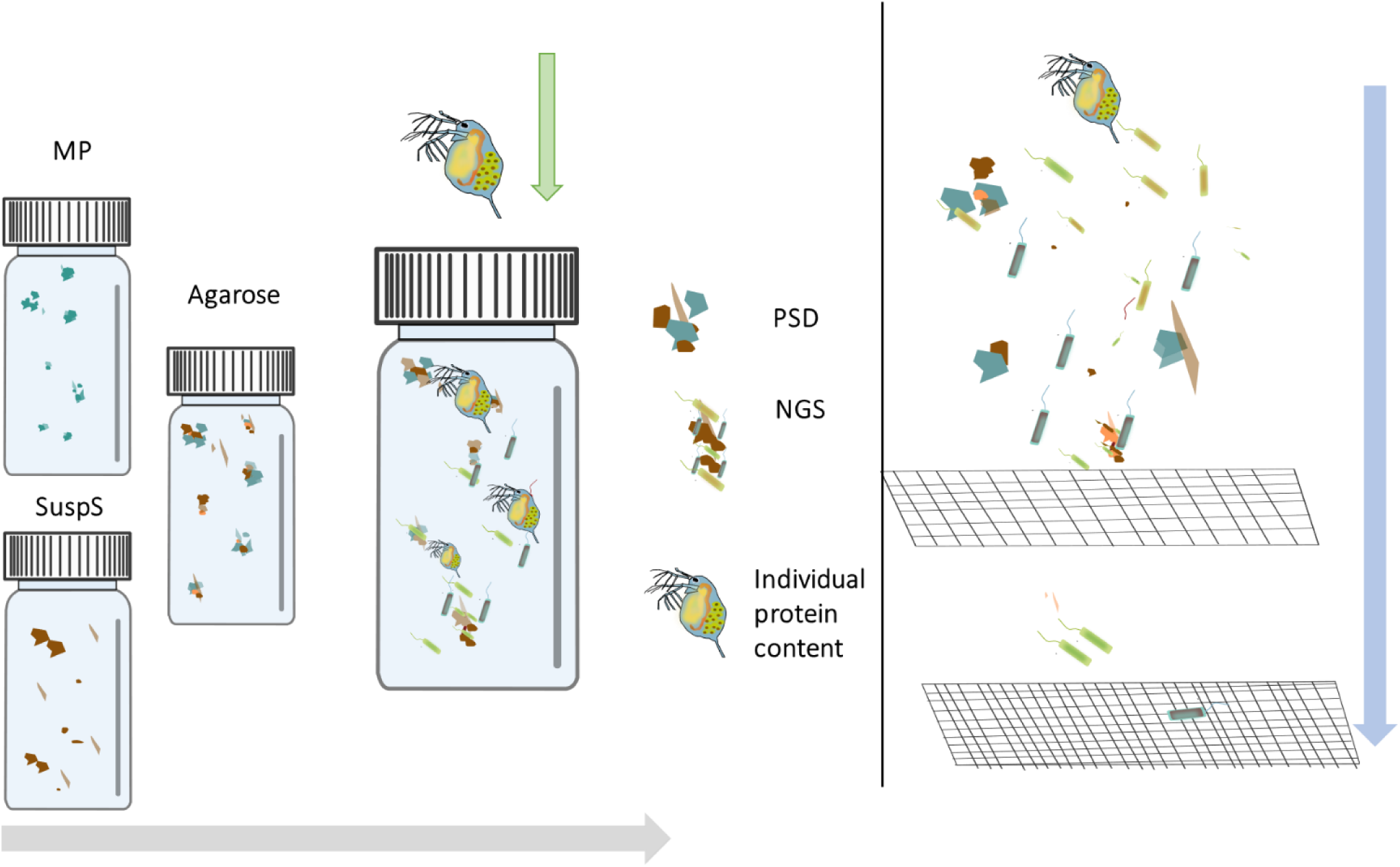
A: Experimental factors and measured endpoints. (A) The daphnids were added to vials containing particle aggregates that were generated using MP and reference particles, agarose, and M7 media. The daphnids served as donors to inoculate the system with their microbiota. After 72 h, the daphnids were collected for protein measurements. (B) The remaining exposure media with aggregates and bacteria was sequentially filtered, first with a 3-µm filter to collect the aggregates and then with a 0.2-µm filter to collect non-adherent bacteria. The filters were frozen at -80°C.

At the end of the experiment, the mortalities were recorded, and the surviving daphnids were snap-frozen (-80 ^◦^C). These individuals were used for protein content measurements, and individual protein content was used as an endpoint of *Daphnia* response to the exposure. The particle suspension was sequentially filtered using first a 3-µm filter (25 mm MERCK) to collect particle-associated bacteria (hereafter referred to as *biofilm*) and passing the filtrate onto a 0.2-µm filter (Merck) to collect non-adherent cells; all filters were snap-frozen and stored at -80^◦^C. These filter samples were used for DNA extraction.

As blanks, all media, particle stocks, and UV-thermo-treated filters were used to extract DNA and assess procedural contamination; the DNA extraction was conducted using the methods described in section 2.3.2. Although detectable DNA concentrations were not recorded in the blank samples, they were used for PCR, yielding no positive amplification. Thus, we assume that all bacteria in both 0.2- and 3-µm fractions originated from the *Daphnia* microbiome. The *Daphnia* microbiome used for the inoculation was assessed using 10 individuals at the start of the experiment pooled into a composite sample and processed in parallel with the bacterial samples, as described in section 2.3.2.

### 2.3 Endpoints

#### 2.3.1 Protein content

As the animals were not feeding during the exposure, their body mass, including protein, was declining due to the metabolics costs; the higher losses were interpreted as worsened body condition. Individual protein content was quantified before and after exposure using the bicinchoninic acid (BCA) protein kit (Thermo Scientific ^TM^, Micro BCA™ Protein Assay Kit). Each individual was homogenised in 150 µl Milli-Q water using a Hand-held homogeniser (BT Lab System). In a microplate, 50 μL of the homogenate (blank) were mixed with 100 μL of distilled water per well. To each well, 150 μL of the BCA reagent were added, and the plate was incubated at 37°C for 2 h. The absorbance was measured at 562 nm in a plate reader (FLUOstar Optima) using a standard curve with bovine serum albumin.

#### 2.3.2 DNA extraction and 16S rRNA gene amplification

DNA was extracted from each sample using 10% Chelex suspension (Straughan and Lehman, 2000). The DNA quality and quantity were first screened using Nanodrop (NanoPhotometer; Implen), followed by a PCR (StepOne plus real-time PCR systems; Applied Biosystems) to target V3-V4 region with primers 341F 5′-CCTACGGGNGGCWGCAG-3′ and 805R 5′-GGACTACHVGGGTWTCTAAT-3′ (Herlemann et al., 2011). The Master mix (25 µl per reaction) contained 5 μl of each primer (1 μM), 10 μl of Phusion Buffer (Invitrogen), 0.25 µl of Phusion polymerase, 0.05 µl of dNTPs (10 mM, Invitrogen) and 2.5 μl of DNA template. The amplification protocol consisted of an initial denaturation at 98°C for 30 secs followed by 35 cycles of 10 sec at 98°C, 30 sec at 58°C and 72°C, and a final extension step (72°C for 10 min). The PCR products were cleaned with the AMPure kit (Qiagen).

#### 2.3.3 Library preparation

Denaturation was carried out using 0.2 N NaOH and sequenced with the MiSeq Illumina system (2 × 300 bp paired-end). Demultiplexing and removing indexes, primers, and tags were handled by the Illumina software (v 2.6.2.3) following the Illumina protocol. Low-quality sequences were detected and removed using DADA2 version 1.6 for R statistical software (v3.4.2, R Core Team, 2018). The pipeline further includes error estimation and de-replication. Paired-end reads to obtain amplicon sequence variants (ASVs) were merged and cleaned by filtering the Phred quality score (Q20 and Q30) to generate data for operational taxonomic units (OTUs) clustering and taxonomy analysis. Sequences were cut 60 and 100 bp downstream of the forwards and reverse primer, respectively, generating a pb fragment for the downstream statistical analyses. Finally, chimeric and ambiguous sequences were removed, and the resulting sequences were matched with the SILVA taxonomy database (138.1). Finally, the eukaryotic and chloroplast ASV were removed. The subsequent statistical analysis was performed with the *Phyloseq* R-module v 1.34.0. All sequences are available at …

### 2.4 Data analysis

The data analysis was organised based on the three primary hypotheses (Table 3).

**Table 2.** Hypotheses addressed in the study.

| # | Hypothesis |
| --- | --- |
| 1 | Bacteria in both size fractions (0.2-µm and 3-µm) originate from the *Daphnia* microbiome. |
| 2 | Experimental factors and PSD have specific effects on each size fraction: |
|  | 2a: Agarose influences primarily the non-adhering bacteria serving as a carbon source and changing the viscosity of the media. |
|  | 2b: Parameters related to solid particles (both their total concentration of and MP contribution) influence primarily particle-associated bacteria (biofilm) because these factors change aggregate size and topology (Reichelt and Gorokhova, submitted). |
| 3 | Responses of *Daphnia* to the exposure conditions are primarily driven by PSD (Motiei et al., 2021) and bacteria in the system (Gorokhova et al., 2021). |

**Table 3.** GLM output for the alpha diversity indices (Shannon-Wiener and Fisher’s alpha) and the experimental factors in (A) 0.2-µm fraction (non-adhering cells) and (B) 3-µm fraction (biofilms). The most parsimonious model was identified as a best-fit model with the fewest number of predictors using AIC. Significant p values are in boldface.

|  | Level of effects | Estimate | Standard Error | Wald statistic | P value |
| --- | --- | --- | --- | --- | --- |
| **A. 0.2-μm fraction (non-adhering cells)** |  |  |  |  |  |
| *Fishers' alpha* |  |  |  |  |  |
| Skewness |  | 0.2646 | 0.1291 | 4.202 | **0.0404** |
| Agarose | 0 | -0.0950 | 0.0382 | 6.190 | **0.0128** |
| **B. 3-μm fraction (biofilm)** |  |  |  |  |  |
| *Shannon-Wiener- index* |  |  |  |  |  |
| Agarose | 0 | -0.04302 | 0.02361 | 3.318 | 0.0685 |
| MP>1 | 1 | -0.077132 | 0.0262 | 8.647 | **0.00327** |
| Mode >1 | 1 | -0.06552 | 0.0236 | 7.661 | **0.00564** |
| *Fishers' alpha* |  |  |  |  |  |
| Agarose | 0 | 0.02111 | 0.03901 | 0.293 | 0.58851 |
| SS <1 | 0 | 0.1158 | 0.03723 | 9.678 | **0.0018** |
| MP >1 | 1 | -0.162997 | 0.04463 | 13.334 | **0.00026** |
| Mode >1 | 1 | -0.06587 | 0.03848 | 2.930 | **0.0869** |
| Agarose × SS <1 | 1 | 0.10557 | 0.03850 | 7.5419 | **0.0061** |

#### 2.4.1 Bioinformatics

Illumina MiSeq sequencing resulted in a total of 5369870 reads among all samples. The average was 42958. The OTU number was 8705 with 3088 for ≥ 2 counts. Sequences were filtered to a minimum count of 4 and a prevalence of 20%. Additionally, a low variance filter was applied to remove taxa with a minor variation throughout the samples at a threshold of 10%, and all data were log-centered. No rarefaction was needed due to similar sequencing depth among the samples.

Shannon-Wiener and Fisher’s alpha were identified as two complementary alpha diversity indices (Figure S4; Supplementary Information) using Principal Component Analysis. These indices were calculated for each size fraction of bacteria using non-filtered data at the ASU level (R package *Phyloseq*, version 1.16.2).

Differential taxon relative abundances were calculated using DESeq2 as implemented in the Marker Data Profiling pipeline from MicrobiomeAnalyst (Dhariwal et al., 2017) with default settings on 25 March 2023 to identify taxa significantly related to the experimental factors and PSD. DESeq2 uses count data to detect differentially expressed taxa using negative binomial generalised linear models. It provides estimates of dispersion and logarithmic fold change (log2FC; log2 fold-change). In this analysis, we used SS, MP%, and Agarose as well as binarised PSD metrics (D_10_, D_50_, D_90_, Kurtosis, Skewness, and the number of modes in the particle size spectra) as predictors (Table 1). In addition, SS and MP% were tested as both 3-level factors and as binarised categorical variables in the search for the most significant model (Table S3). When both 3-level and 2-level factors generated from the same continuous variable produced similar results, the model with the highest Log2FC was selected for the presentation.

#### 2.4.2 Effects of the experimental factors and PSD on bacterial communities

Generalised linear models (GLM) were used to complement the DESeq2 results and analyse relationships among the variables in combination, thus putting them in a broader context. GLMs with log-link function were applied to identify significant drivers (R package MASS, version 7.3.53; Statistica, Tibco Software) of each response variable (i.e., Shannon-Wiener and Fisher’s alpha indices and relative abundances of the impacted taxa) with SS, MP%, and Agarose as well as PSD metrics (D_10_, D_50_, D_90_, Kurtosis, Skewness and the number of modes in the particle size spectra) as predictors (Table 1). The binarised predictor variables used for DESeq2 were also tested in these models because the responses to the continuous preditors were not monotonous (Figure S6). The most parsimonious model was selected as the best-fit model with the fewest predictors using the Model Building Module in Statistica and the Akaike Information Criterium (AIC).

#### 2.4.3 Aggregation and microbial communities as predictors of *Daphnia* responses

GLM was also used to assess the effects of the experimental factors (Table 1), PSD (D_10_, D_50_, D_90_, skewness, and kurtosis), and bacterial diversity (Shannon-Wiener and Fishers’ alpha indices) on *Daphnia* response variables (individual protein content and mortality). Untransformed data, a log-link function, and a normal error structure were used. Finally, we applied DESeq2 to identify taxa with up- or downregulated abundances that were associated with adverse responses in *Daphnia*, i.e., high mortality (>10%; a conventionally accepted value for control mortality in acute toxicity testing) and low individual protein content (i.e., values below the median for the entire data set).

## 3. Results

### 3.1 Bacterial communities in the exposure system

#### 3.1.1 Library size

The raw ASV number ranged from 28857-96397 (all samples), 29199-66257 (0.2-um fraction), and 29430-73238 (3-um fraction) (Figure S1), whereas *Daphnia* microbiome contained 57563 ASVs. All samples reached saturation level; therefore, the sequencing depth was considered sufficient (Figure S2). Following all filtering steps, 273 features were retained for the analysis.

#### 3.1.2 Similarity between the *Daphnia magna* microbiome and bacteria communities in the test system

*Daphnia* microbiome and both size fractions were dominated by Flavobacteriaceae, Sphingobacteriaceae, Comamonadaceae, unclassified_Blfdi19 (Myxococcota), and Cyclobacteriaceae (Figure 2, Figure S3). Bacterial genera in the exposure system were similar to those found in the *Daphnia* microbiome, with 57% overlap between all three communities and 5% being unique to the *Daphnia* microbiome (Figure 3). The ASV number was similar between the 0.2-µm and 3-µm fractions, 236 and 213, respectively, and 16% of them were shared between these size fractions.

**Figure 2.**
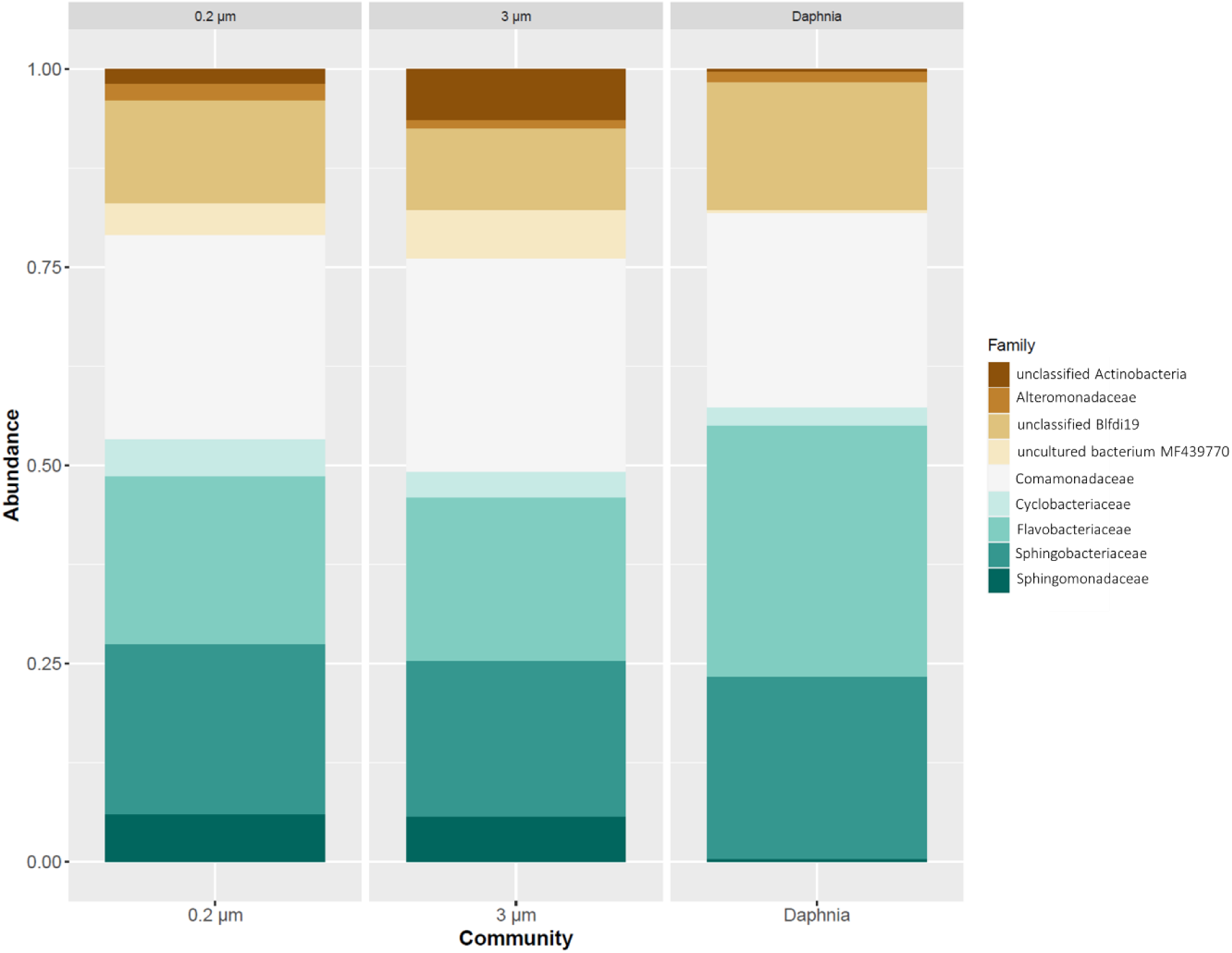
Relative abundances in 0.2-µm fraction (non-adhering cells), 3-µm fraction (biofilm), and the Daphnia microbiome. Data are presented at the family level when possible. Unclassified taxa are referred to by the closest taxonomic category.

**Figure 3.**
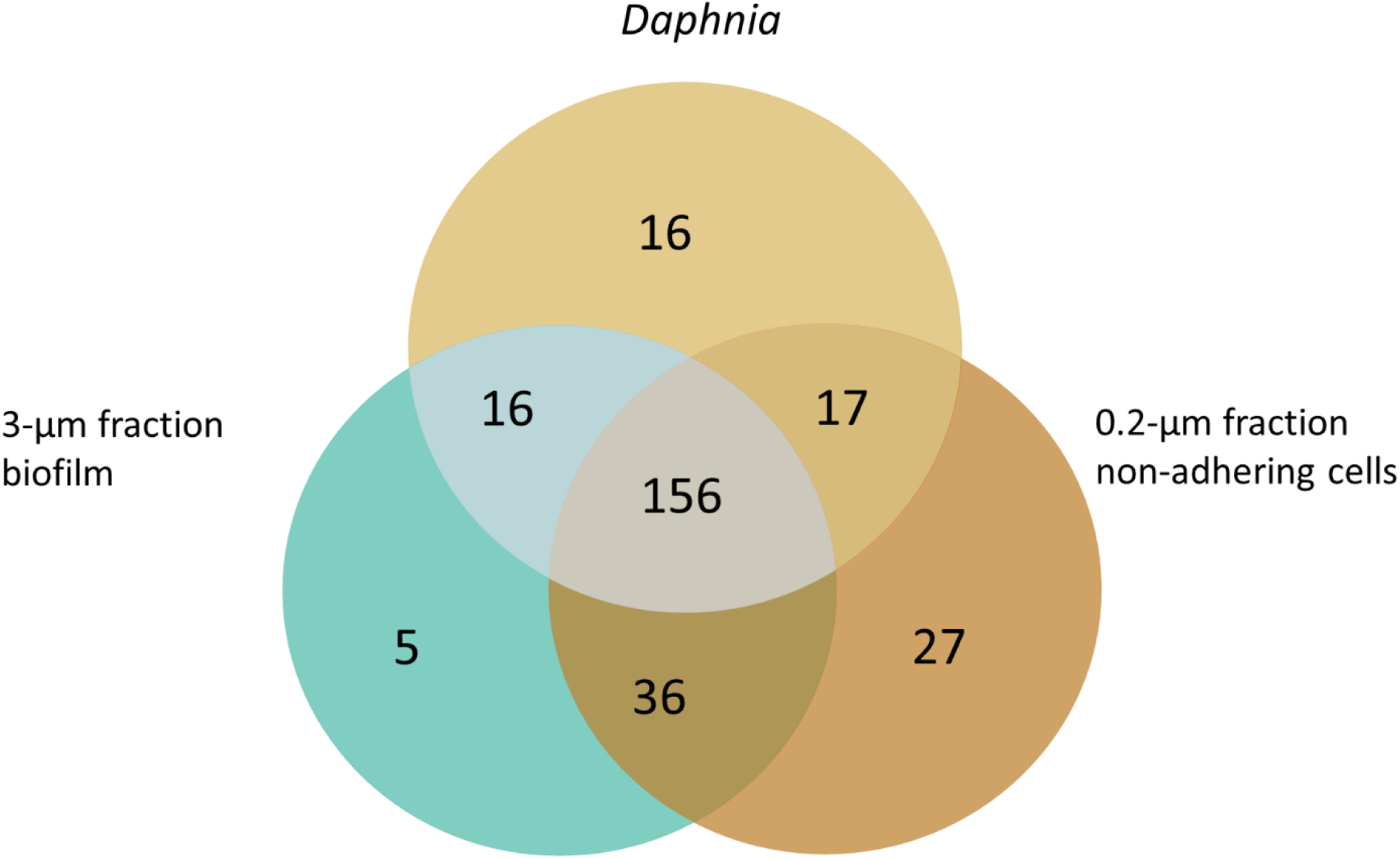
Venn diagram for ASV diversity in the test communities. All communities share 156 ASVs.

**Figure 4.**
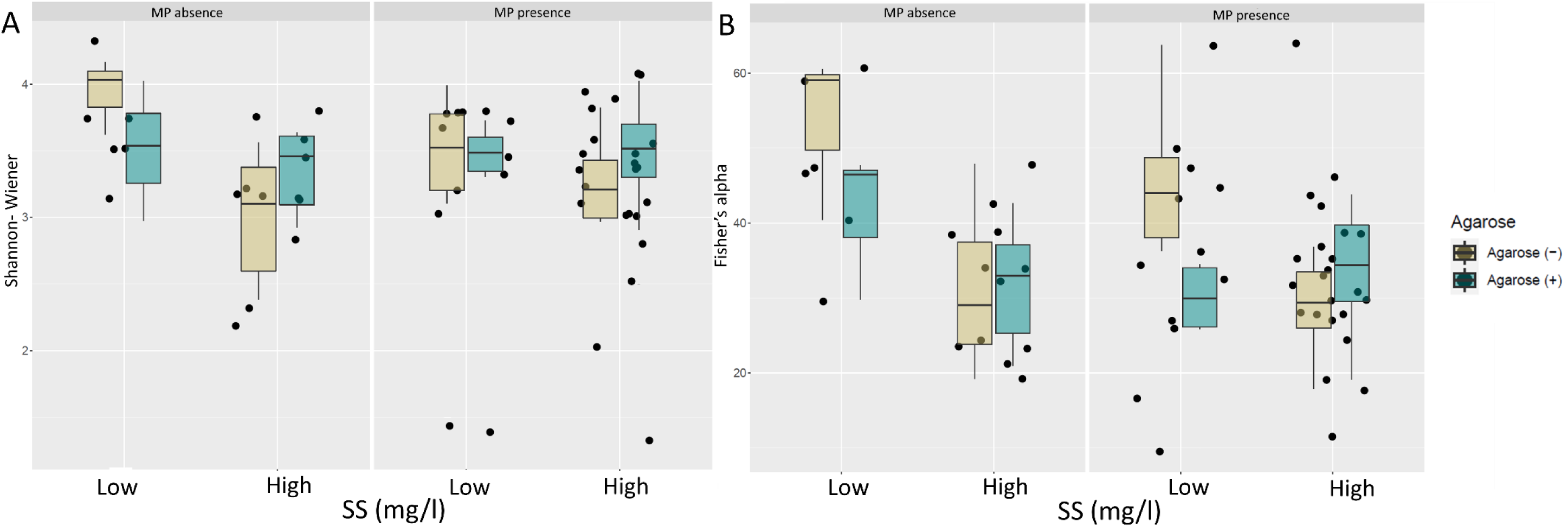
Alpha diversity is assayed as (A) Shannon-Wiener index and (B) Fisher’s alpha. The x-axis shows the suspended solids as a binary variable low/high). The box shows 50% of the data confined by the first and third quartile, and the whiskers show a 95% confidence interval. The color code for the agarose treatment indicates presence/absence.

**Figure 5.**
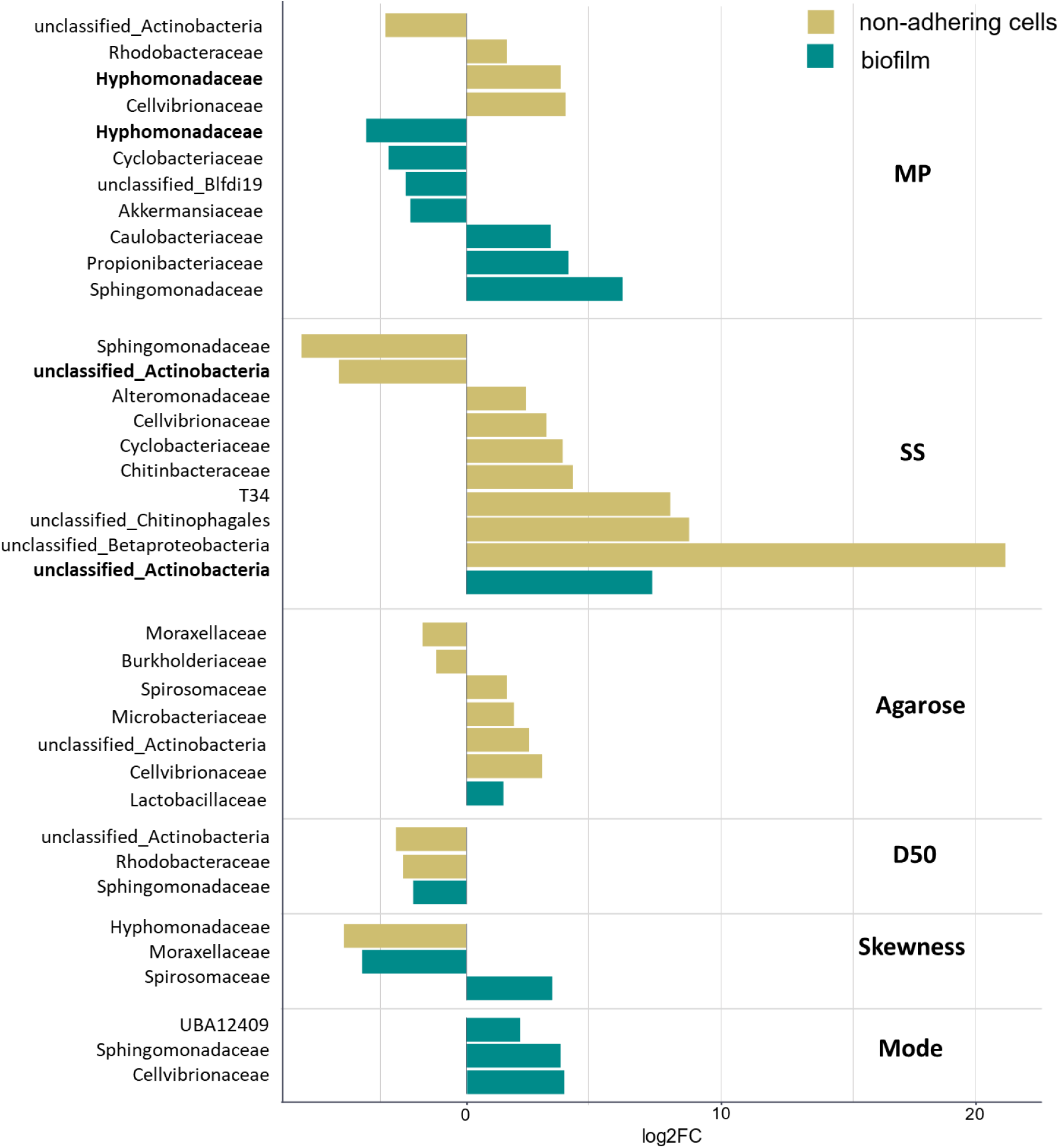
DESeq2 results for the treatment effects and PSD metrics the differential abundance of bacteria at the family level in the biofilm (3-um fraction, green) and non-adhering cell samples (0.2-um fraction, yellow).

The *Daphnia* microbiome was comprised mainly of four families making up 90% of the relative abundance: Flavobacteriaceae (26%), Comamonadaceae (25%), Sphingobacteriaceae (23%), and unclassified Bldfi19 (18%). The most frequent genera were Pedobacter, Flavobacterium, Aquabacterium, and Algoriphagus (Figure 2).

The dominant families in the seeded communities were Comamonadaceae (27%), Sphingobacteriaceae (22-23%), Flavobacteriaceae (22%), uncultured bacteria MF439770 (5-11%), unclassified Blfdi (3-6%), Actinobacterium (2-6%), and Sphingomonadaceae (5%) (Figure 2). Hyphomonadaceae family was significantly less frequent in biofilms than in the non-adhering community (DESeq2: FDR < 0.05). The most frequent genera within the dominant families were Variovorax, Pedobacter, Flavobacterium, Aquabacterium, Cyclobacteriaceae and unclassified Actinobacterium.

### 3.2 Effects of experimental factors and PSD Parameters on bacterial communities

#### 3.2.1 Effects on the alpha diversity Indices

The data exploration suggested that the diversity indices and their responses to the experimental factors differed between the bacteria size fractions (0.2-µm vs 3-µm; Figure 3, Table S4, Table S5, Figure S5). Therefore, the GLMs evaluating these responses for each size fraction explored all experimental factors (SS, MP%, Agarose, and selected PSD parameters [number of modes, D_50_, D_10_, D_90_, skewness, and Kurtosis]) and 2-way interactions as predictors (Figure S5).

No significant model was identified for the 0.2-µm fraction (i.e., non-adhering cells) with the Shannon-Wiener index as a response variable. In contrast, we found negative effects of the agarose addition and PSD skewness for Fisher’s alpha index, implying that PSD asymmetry with a prevalence of smaller particles and a lower proportion of large aggregates increases bacterial diversity. Neither SS nor MP% were identified as significant predictors.

For the 3-µm (i.e., biofilms), both Shannon-Wiener and Fisher’s alpha indices were affected by at least one PSD parameter and the experimental factors (Figure 3). Significant negative effects of MP% and PSD multimodality were found for Shannon-Wiener diversity; moreover, agarose addition had a negative, marginally significant effect and was retained in the model. Similarly, Fisher’s alpha was lower when MP were more than 1% and PSD was multimodal. Finally, a significant Agarose × SS interaction was detected, implying that agarose addition increased Fisher’s alpha at SS levels exceeding 1 mg/l.

#### 3.2.2. Effects of the experimental factors and PSD parameters on the differential abundance of bacterial taxa

The DESeq2-tool was used to identify responsive taxa for each experimental factor (Figure 6, Table S5). In total, 11 families in both size fractions were affected by MP exposure, with approximately the same probability of up- and downregulation of the relative abundances (Figure 6 A). In the 0.2-µm fraction, Rhodobacteraceae, Cellvibrionaceae, and Hyphomonadaceae were upregulated, whereas unclassified Actinobacteria were downregulated. In the 3-µm fraction, four families (Hyphomonadaceae, Cyclobacteriaceae, unclassified Blfdi19, and Akkemansiaceae) were downregulated, and three (Caulobacteraceae, Propionibacteriaceae, and Sphingomonadaceae) were upregulated. Thus, Hyphomonadaceae were affected by MP in both communities, albeit with the opposite effects.

**Figure 6.**
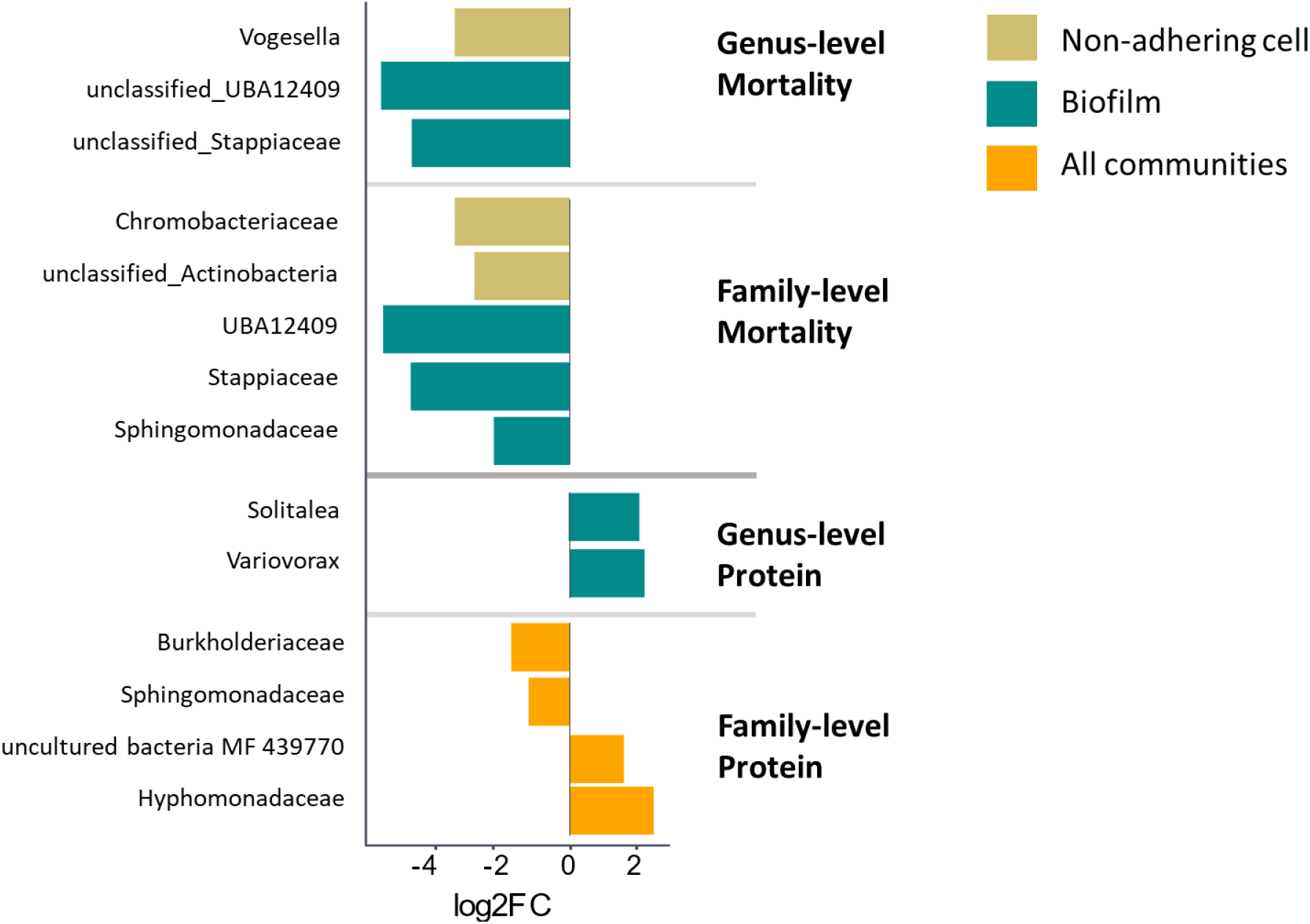
DESeq2 results for the Daphnia mortality and individual protein content on the differential abundance of bacteria at the family- and genus levels in the biofilm (3-um fraction, green), non-adhering cell samples (0.2-um fraction, yellow) and all communities pooled (orange). A high log2FC is associated with low mortality and high protein.

The high levels of the suspended solids (>1 mg/l) had stimulating effects on the 0.2-µm fraction, where seven families were mostly upregulated (2.3 to 21.2 Log2FC), with the greatest fold-change compared to the other experimental factors (Figure 6 B). The most influenced taxa were unclassified Betaproteobacteria, unclassified Chitinophagales, and T34. Notably, unclassified Actinobacteria were suppressed by SS and MP% in the 0.2 µm fraction and stimulated by SS in the 3-µm fraction. Thus, MP% was the most influential experimental factor, with the most profound effects on the biofilm communities. Moreover, MP% and SS had opposite effects on some bacteria, i.e., Sphingomonadaceae and Cyclobacteriaceae. The former was upregulated by MP% and downregulated by SS; the latter was upregulated by SS and downregulated by MP%.

Like the suspended solids, agarose mainly affected the 0.2-µm fraction, stimulating Spriosomaceae, Microbacteriaceae, unclassified Actinobacteria, and Cellvibrionaceae and suppressing Moraxellaceae and Burkholderiaceae. In the 3-µm fraction, only Lactobacilliaceae were significantly upregulated in the agarose treatments (Figure 6 C).

The aggregate size and diversity also affected the relative abundance of some bacteria in both size fractions (-4.8 to 3.7 Log2FC). The significant upregulation in biofilms (Sphingomonadaceae, Cellvibrionaceae, and UBA12409) were observed in multimodal aggregates, whereas skewness had both upregulating (Spirosomaceae) and downregulating (Hyphomonadaceae and Moraxellaceae) effects. The non-adhering Actinobacteria (unclassified), Rhodobacteriaceae and Hyphomonadaceae decreased with decreasing the aggregate size in the system.

#### 3.2.3. Effects of the experimental factors and PSD on biofilms

The relative abundances of the biofilm families responding to any of the experimental treatments were used as response variables in GLMs with experimental factors (SS, MP%, and Agarose) and PSD parameters (D_10_, D_50_, D_90_, Skewness, Kurtosis, Number of modes, Mode) as predictors (Table S3). At least one PSD metric was a significant predictor in all best-fit GLMs for the biofilm families affected (as identified by DESeq2). Although the fold-change values for PSD parameters were not the highest among all factors tested by DESeq2, the GLM output suggested that the PSD metrics were among the strongest predictors when combined with other experimental factors (Table S7). The most frequent PSD predictors were kurtosis, skewness, and the number of modes (i.e., PSD multimodality). Each of these predictors was associated with both positive and negative effects on the relative abundances in different families. For example, Cyclobacteriaceae, Blfdi19, and Lactobacillaceae were positively affected by skewness, whereas Actinobacteria were affected negatively. Thus, the latter were more abundant when PSD was shifted toward the larger aggregates and the former preferred suspensions with the bulk of the particles being small. Also, some bacteria (e.g., Actinobacteria, Blfdi19) were associated with flat-topped PSD (platykurtic), whereas others (Cyclobacteriaceae and Caulobacteriaceae) preferred narrow peaks (leptokurtic), characteristic of higher aggregate size similarity. In line with the kurtosis and skewness effects, multimodality was associated with increased Blfdi19 and decreased Cyclobacteriaceae and Lactobacillaceae. Among the PSD metrics, the diversity of the aggregates manifested as an increased number of modes and decreased skewness were most commonly associated with the increased relative abundances of the responsive families in biofilms. Thus, the GLMs implicate aggregate size and topology as the primary drivers of the taxa affected by the exposure.

Exposure to MP had significant effects in 4 out of 7 models, with positive effects on the relative abundance of Caulobacteriaceae and Sphingomonadaceae and adverse effects on Actinobacteria and Blfdi19. Agarose was also a significant negative predictor in 4 out of 7 models, strongly affecting Caulobacteriaceae and Akkermansiaceae. In combination with at least one PSD-related variable, suspended solid level was a significant predictor in three models, with adverse effects on Cyclobacteriaceae and Actinobacteria and positive on Caulobacteriaceae.

### 3.3. Effects of the experimental factors, PSD, and bacterial communities on *Daphnia*

Experimental factors (MP%, SS, Agarose) together with PSD parameters (D_50_, D_90_, D_10_, mode, kurtosis, and skewness) and bacterial diversity indices (Shannon-Wiener- and Fisher’s alpha diversity indices) were tested as predictors in a GLM with individual protein content as the *Daphnia* response. Agarose and Fisher’s alpha were significantly negative predictors, while median aggregate size (D_50_) significantly positively affected daphnid protein content (Table 4).

**Table 4.** Best-fit GLM for the relationship between the individual protein content of Daphnia magna and the retained predictors (agarose, Fisher’s alpha for bacteria diversity of biofilm, and median aggregate size).

|  | Estimate | Std. Error | P-value |
| --- | --- | --- | --- |
| Agarose | -0.07 | 0.03 | **0.035** |
| Fisher's alpha, biofilm | -0.00429 | 0.002 | **0.035** |
| $D_{50}$ | 0.00323 | 0.0015 | **0.032** |

Further, DESeq2 identified families in biofilm communities associated with elevated (above 10%) mortality and reduced (below the median value for the group) individual protein content. Daphnids survived better when unclassified UBA12409, Stappiaceae, Sphingomonadaceae, Chromobacteriaceae and unclassified Actinobacteria were more prevalent in the biofilms (Figure 6). Concerning the taxa associated with the protein content, Hyphomonadaceae and uncultured bacteria MF 439770 were upregulated in the high-protein daphnids, whereas Sphingomonadaceae and Burholderiaceae were downregulated (Figure 6); these differences were observed when both biofilms and non-adhering cells were considered. Also, higher frequency of the genera *Solitalea* and *Variovorax* in biofilms improved the protein content.

## 4. Discussion

### 4.1. Effects of the experimental factors on the assay microbiome

The role of microorganisms in effect studies with particulate materials, such as microplastics, is often overlooked. We examined how the microbial communities associated with *Daphnia magna* inoculate a test system containing a mixture of clay and microplastics. We focused on the particle-associated biofilms and non-adherent cells to understand if/how (1) experimental factors affect their diversity and composition and (2) daphnids respond to these bacteria in the test system. We found that the experimental factors influenced the non-adherent cells and biofilms differently, with total suspended solids and DOM primarily affecting non-adhering cells and microplastics impacting biofilms. Moreover, the aggregate size and topology strongly influenced bacterial diversity and the abundance of specific families in the test system. Additionally, we observed adverse effects on survival and individual protein content in *Daphnia*, driven by small aggregate size, agarose addition, and high biofilm diversity.

As hypothesised (H1), the microbiota of *Daphna magna* was transferred to all compartments of the test system; bacteria colonised particle surfaces and occurred in the water as free-living cells. Thus, even though the sterile flasks and medium were used and the test particles were bacteria-free (as shown by no amplification of the 16S rRNA gene in the blanks), the experimental system developed a microbiome that originated from the test organism. Therefore, any ecotoxicity test system with a test organism that is not germ-free will have a microbiome as a component interacting with other system components, dissolved and particular matter and test organisms, and potentially contributing to the test outcome (Duperron et al., 2020). This also holds for testing dissolved chemicals, such as pesticides or heavy metals, and potentially harmful particles; however, the biofilms on the test particles are especially important when testing suspended solids, such as nano- and microparticles, because they can influence particle uptake by the test organism (Licht and Bahl, 2019; Maszczyk et al., 2022).

The host and the assay microbiome shared 56% of the total ASVs, with about 5% being unique to the *Daphnia* microbiome, and 1% and 5% ASVs being unique to the non-adhering cells and the biofilms, respectively. Thus, most ASVs in the inoculum were transmitted to the media and established as biofilms or non-adhering cells. Moreover, the unique ASVs found in the biofilms and media were, most likely, rare taxa not readily identifiable in the donor communities. In contrast, the unique ASVs in the *Daphnia* microbiome represented taxa that were not readily adaptable to the environment outside of the host body. Therefore, the composition of the inoculum, selection in different microhabitats of the test system and migration dynamics between the compartments (i.e., media ↔ biofilm) during the exposure affected the microbiome of the test system.

The bacterial communities of the two size fractions shared a majority of taxa, with some taxonomic differences between them. For example, Hyphomonadaceae family was significantly more frequent in the non-adhering community than biofilms, which was in line with other reports on their association with particulates (Park et al., 2021). The opposite was found for the unclassified Actinobacteria that were more common in the biofilms and contributed significantly to the differences between the non-adhering cells (1.6%) and biofilms (7%) (Figure 2), supporting observations of attached Actinobacteria in the detrital and sediment particles, high OTU richness and phylogenetic diversity in filed populations (Parveen et al., 2011). As expected (Table 2; H2), the effects of the experimental factors varied between the non-adhering and biofilm fractions due to these taxonomic differences and the fact that attached bacteria generally show higher cell-specific growth rates than their free-living counterparts in aquatic ecosystems and respond to different cues (Fandino et al., 2001; Lemarchand et al., 2006).

Agarose addition was the most influential factor for the non-adhering cells, decreasing alpha diversity and affecting six families with mostly upregulating effects. Thus, the hypothesised prevalence of the agarose effects on the non-adhering cells (H2a) was supported. The agarose addition to the media can select for agarose-degrading bacteria that possess the necessary enzymes to break down agarose into simpler sugars, which may explain upregulation of Spirosomaceae, Microbacteriaceae, unclassified Actinobacteria and Cellvibrionaceae. Also, agarose alters the physical properties of the media, such as viscosity or nutrient availability, and contributes to the formation of microhabitats or niches within the media, which may influence the colonisation and persistence of specific bacterial groups.

The influence of the total suspended solid concentrations on the alpha diversity was found for biofilms only; moreover, it was conditional on agarose addition, with decreased Fisher’s alpha in no-agarose/high SS treatments. Overall, SS was a very influential factor on the taxonomic composition of the non-adhering cells, where high SS levels were primarily associated with the upregulation of seven families that was exceptionally high for some taxa (e.g., >20-fold for unclassified Betaproteobacteria [Figure 6] and >23 for actinobacterial Aurantimicrobium; data not shown). Moreover, the magnitude of upregulation was much stronger than downregulation. Thus, as hypothesised, more substantial effects of particulate material on the biofilms compared to the non-adhering cells were partially supported for the total suspended material and demonstrated for microplastics (H2b). The bacterial families upregulated by SS were Alteromonadaceae, Cellvibrionaceae, Cyclobacteriaceae, Chitinbacteraceae, T34, unclassified Chitinphagales and unclassified Betaproteobacteria (non-adhering fraction) and unclassified Actinobacteria (biofilm). Numerous reports support the occurrence of various genera belonging to these families in turbid environments (Chavarria et al., 2021; Dodd et al., 2020).

The biofilm community was significantly affected by an increase in MP above 1% (of the total suspended solids by mass), which emerged as the most influential factor for suppressing the alpha diversity and altering the relative abundances of seven families (Figure 6A). Notably, due to the lower specific gravity of polystyrene compared to kaolin, this threshold would correspond to 1.57% by the particle number. In our aggregates, kaolin was the primary material, with the highest abundance of 10 mg/l and the highest MP contribution of 10%. Therefore, the relative surface of the polymer available for colonisation was small compared to the mineral surface, yet, the MP effects were detectable.

MP upregulated Caulobacteraceae, Propionibacteriaceae, and Sphingomonadaceae, while downregulated Hyphomonadaceae, Cyclobacteriaceae, unclassified Blfdi, and Akkermansiaceae. Thus, families responding to both MP and SS were Sphingomonadaceae (upregulated by MP and downregulated by SS) and Cyclobacteriaceae (downregulated by MP and upregulated by SS) (Figure S6). Hence, MP and SS had opposite effects on these bacteria, suggesting that the net effect of the mixture would vary depending on the relative contribution of each particle type in the aggregates.

Other studies reported similar effects of MP (Miao et al., 2019), although not in mixed aggregates. Also, early-stage colonisers have been reported to be significantly different between MP and natural surfaces as well as the surrounding water (Di Pippo et al., 2020). Moreover, in line with our findings, PS particles at 10 mg/l reduced alpha diversity in freshwater biofilms (Miao et al., 2019) and - at 100 µg/l - the gut microbiome of medaka (Yan et al., 2020). Yet, the opposite was found in the biofilms on the Atlantic coast and mesocosms running with the Baltic Sea bacterioplankton, where alpha diversity was higher on plastic compared to the surrounding water (Frère et al., 2018) and natural surfaces (Ogonowski et al., 2018). What is also relevant for experimental design with MP and other particles, anthropogenic or natural, is that bacterial communities associated with e.g., sand, chitin and cellulose foster communities which are characterised by different dominating species indicating that non-polymeric particles can also selectively enrich bacterial assemblages (López-Pérez et al., 2016).

### 4.2. PSD as the primary driver of assay microbiome

By incorporating PSD metrics as predictors, we gained new insights into how the diversity and relative abundance of bacteria responded to the exposure conditions. The regression models identifying the most influential predictors supported the hypothesised role of aggregation as the main driver of the biological responses to particle exposure (H3), with both aggregate size (D_50_, D_10,_ and D_90_) and topology (skewness, kurtosis, and the number of modes) being involved (Table S7). Of all candidate predictors, these metrics were selected by the best-fit models more frequently than the suspended solid concentration (or, alternatively, its binarised counterpart), suggesting that PSD was the proximate driver of the changes in bacterial diversity and community composition observed in the experimental system.

In particular, the increased proportion of large and complex aggregates manifested as an increased number of modes resulted in a smaller total surface area available for colonisation and decreased Shannon-Wiener-diversity index of the biofilms (Table3). Further, Actinobacteria, for example, were associated with some skewness toward larger aggregates with platykurtic distribution, whereas Lactobacilliaceae and Cyclobacteriaceae were more abundant when PSD was highly skewed to the small-sized aggregates (Table S7). Thus, this approach allowed us to demonstrate the importance of particle aggregation analysis in predicting biofilm community structure.

As shown earlier (Reichelt and Gorokhova, submitted), microplastics have direct effects on PSD; however, the best-fit models suggest that they also affected particle-associated bacteria (negatively: Actinobacteria and unclassified Blfdi19; positively: Caulobacteraceae and Sphingomonadaceae). The same is true for agarose, which on the one hand, stimulated aggregation (Reichelt and Gorokhova, submitted) but, on the other hand, suppressed Akkermansiaceae, unclassified Blfdi19, Caulobacteraceae and Lactobacillaceae in biofilms. Therefore, identifying the direct and indirect effects of the test particles on the biofilms and non-adhering bacteria in an exposure system requires monitoring PSD, which plays a critical role in shaping microbial communities.

### 4.3. Aggregation, DOM and microbial communities affect *Daphnia* response

Our findings that microbial communities, especially biofilms, are crucial for animal responses are supported by other studies showing that bacteria adding nutrition (Amariei et al., 2022), fostering pathogenic or rare taxa (Kim et al., 2023), and altering PSD (Rogers et al., 2020) can change exposure conditions and contribute to the variability of the results. We found that particle aggregation, DOM and biofilm formation act in concert to modify the exposure: aggregation reduces the availability of suspended particles in the water, decreasing the encounter rate for daphnids, while biofilm and DOM enhance the aggregation and select for specific taxa that can affect the animal. The net outcome of these processes can have positive and negative implications for the test organisms.

The higher protein content in *Daphnia* was positively associated with aggregate size (D_50_), albeit this effect was of lower magnitude than those of agarose and bacterial alpha diversity. No direct effects of MP and SS were identified (Table 4). The observed better body condition in the animals exposed to larger aggregates was most likely related to the lower energy losses due to filtering activity. At high particle number concentrations, daphnids encounter more particles within a given volume of water and adjust filtering activity to exert more energy for collecting these particles. Also, small particles have the potential to clog the feeding appendages of *Daphnia* and can negatively affect the hydrodynamic properties, further exacerbating the energy losses experienced by the animal.

The adverse effects of agarose on *Daphnia* protein content can be attributed to energy losses caused by the higher viscosity of agarose-containing media. The increased viscosity results in greater resistance for the swimming appendages, leading to additional energy expenditure and potentially impacting the overall body condition (Loiterton et al., 2004).

The presence of high-diversity biofilm communities was found (and expected) to have adverse effects, as biofilms may harbor microorganisms that are less common in the *Daphnia* microbiome. When these taxa become more abundant within the biofilm, the risk of negative interactions with the host organism through ingestion increases, potentially compromising the overall well-being of the biofilm consumer. We observed a high protein content associated with upregulated Hyphomonadaceae and uncultured bacteria MF 439770. Also, the upregulated genera *Solitalea* and *Variovorax* represent a healthy *Daphnia* microbiome (Qi et al., 2009), explaining their positive association with body condition. Conversely, Sphingomonadaceae and Burkholderiaceae were associated with low protein content. Burkholderiaceae contain many known pathogens, and their relative abundances in the *Daphnia* microbiome were low, possibly suppressed by a healthy microbiome. However, these bacteria were overrepresented in the exposure system, and one can speculate that the increased frequency of Burkholderiaceae in the system may have compromised daphnid health.

## Conclusion

The complex nature of MP, combined with its interactions with the environmental matrix, makes it challenging to trace the specific pathways because both indirect and direct effects can occur, often leading to contrasting outcomes. We demonstrated that test particles aggregate in the exposure system, and this aggregation is one of the main drivers of the biological responses. In addition, the animal microbiome introduced to the test system generates diverse bacterial communities forming biofilms on the particle aggregates and occurring as non-adhering cells in the media. Adverse effects on *D. magna* were primarily associated with a small aggregate size and bacterial taxa abundant in biofilms but rare in the microbiome, whereas no direct effects of MP were observed. Understanding aggregation and biofilms underscore the need to account for their interplay in effect studies with particulate materials.

## Supplementary material

**Table S3.** Estimated particle concentration (particle/l) among the treatments based on their mass concentration, density and mean size (Leusch and Ziajahromi, 2021)

| SS | MP% | Kaolin (particle/l) | MP (particle/l) |
| --- | --- | --- | --- |
| 0.1 | 1 | $1.32 \times 10^5$ | $2.08 \times 10^3$ |
| | 10 | $1.20 \times 10^5$ | $2.08 \times 10^4$ |
| 1 | 1 | $1.32 \times 10^6$ | $2.08 \times 10^4$ |
| | 10 | $1.20 \times 10^6$ | $2.08 \times 10^5$ |
| 10 | 1 | $1.32 \times 10^7$ | $2.08 \times 10^5$ |
| | 10 | $1.20 \times 10^7$ | $2.08 \times 10^6$ |

**Table S4.** Chemicals in M7 Daphnia culturing media.

| Constituent | Elendt M7 (mg/l) |
| --- | --- |
| CaCl<sub>2</sub> • 2H<sub>2</sub>O | 293.8 |
| MgSO<sub>4</sub> • 7H<sub>2</sub>O | 123.3 |
| K<sub>2</sub>HPO<sub>4</sub> | 0.184 |
| NaNO<sub>3</sub> | 0.274 |
| NaHCO<sub>3</sub> | 64.8 |
| Na<sub>2</sub>SiO<sub>3</sub> • 9H<sub>2</sub>O | 10.0 |
| H<sub>3</sub>BO<sub>3</sub> | 0.715 |
| KCl | 5.8 |
| LiCl | 0.077 |
| RbCl | 0.018 |
| SrCl<sub>2</sub> • 6H<sub>2</sub>O | 0.038 |
| KI | 0.0033 |
| Na<sub>2</sub>EDTA • 2H<sub>2</sub>O | 0.625 |
| MnCl<sub>2</sub> • 4H<sub>2</sub>O | 0.090 |
| CoCl<sub>2</sub> • 6H<sub>2</sub>O | 0.01 |
| NaMoO<sub>4</sub> • 2H<sub>2</sub>O | 0.016 |
| NaBr | 0.004 |
| CuCl<sub>2</sub> • 2H<sub>2</sub>O | 0.0042 |
| ZnCl<sub>2</sub> | 0.013 |
| Na<sub>2</sub>SeO<sub>3</sub> | 0.0022 |
| NH<sub>4</sub>VO<sub>3</sub> | 0.0006 |
| FeSO<sub>4</sub> • 7H<sub>2</sub>O | 0.0249 |
| KH<sub>2</sub>PO<sub>4</sub> | 0.0143 |
| B12 (cyanocobalamin) | 0.0001 |
| Biotin (d-Biotin) | 0.008 |
| Thiamine (HCl) | 0.075 |

**Figure S6.**
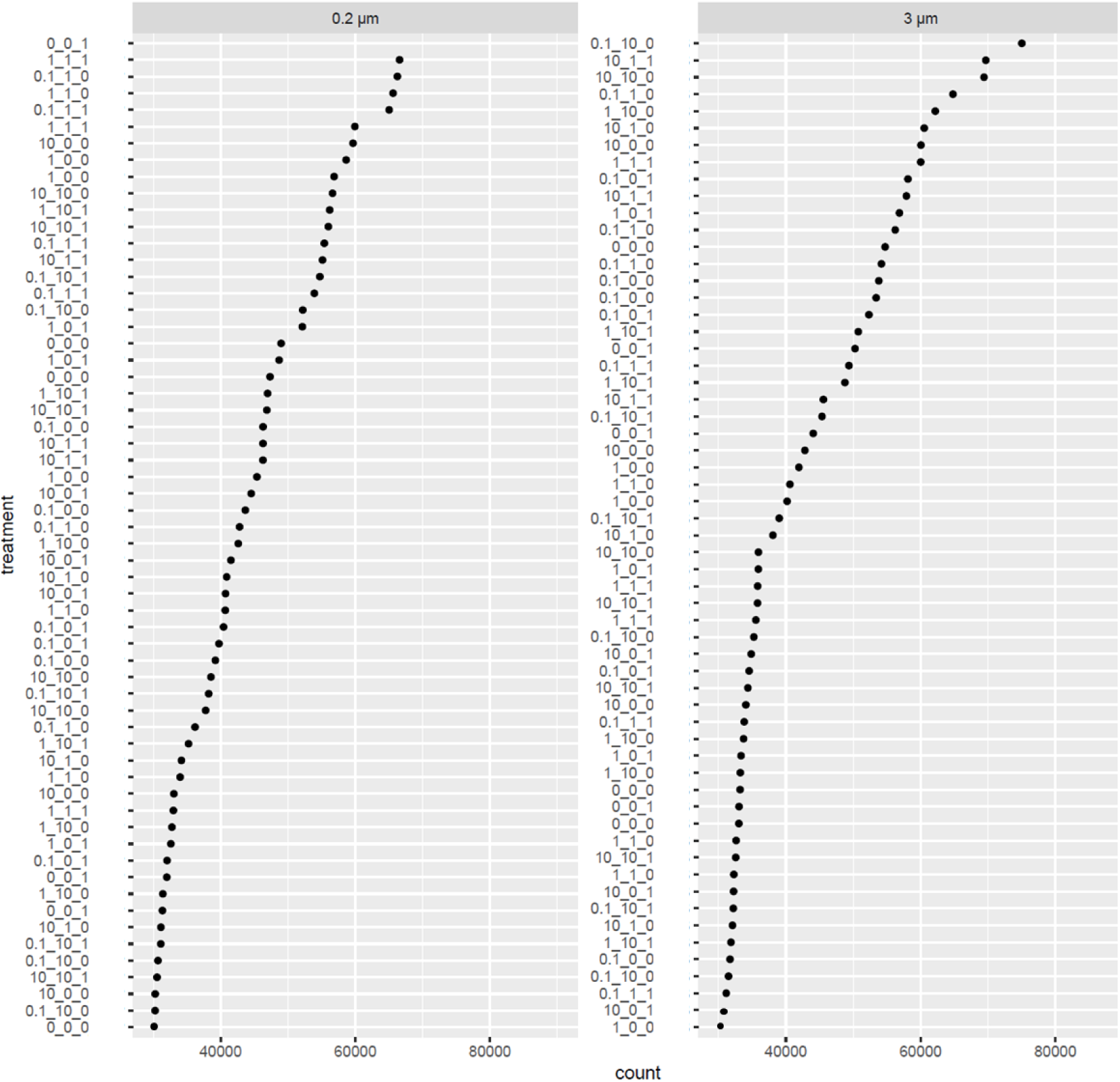
Library size for 0.2-µm and 3-µm bacteria fractions sampled from the experimental units. In the coded sample name, the first number refers to the total SS concentration (0-10 mg/l), the second number - to the MP contribution (0-10%) to the SS, and the third number to the presence/absence of agarose (0: absent, 1: present) in the medium. These raw reads were subjected to filtering and normalization or transformation before the data analysis.

**Figure S7.**
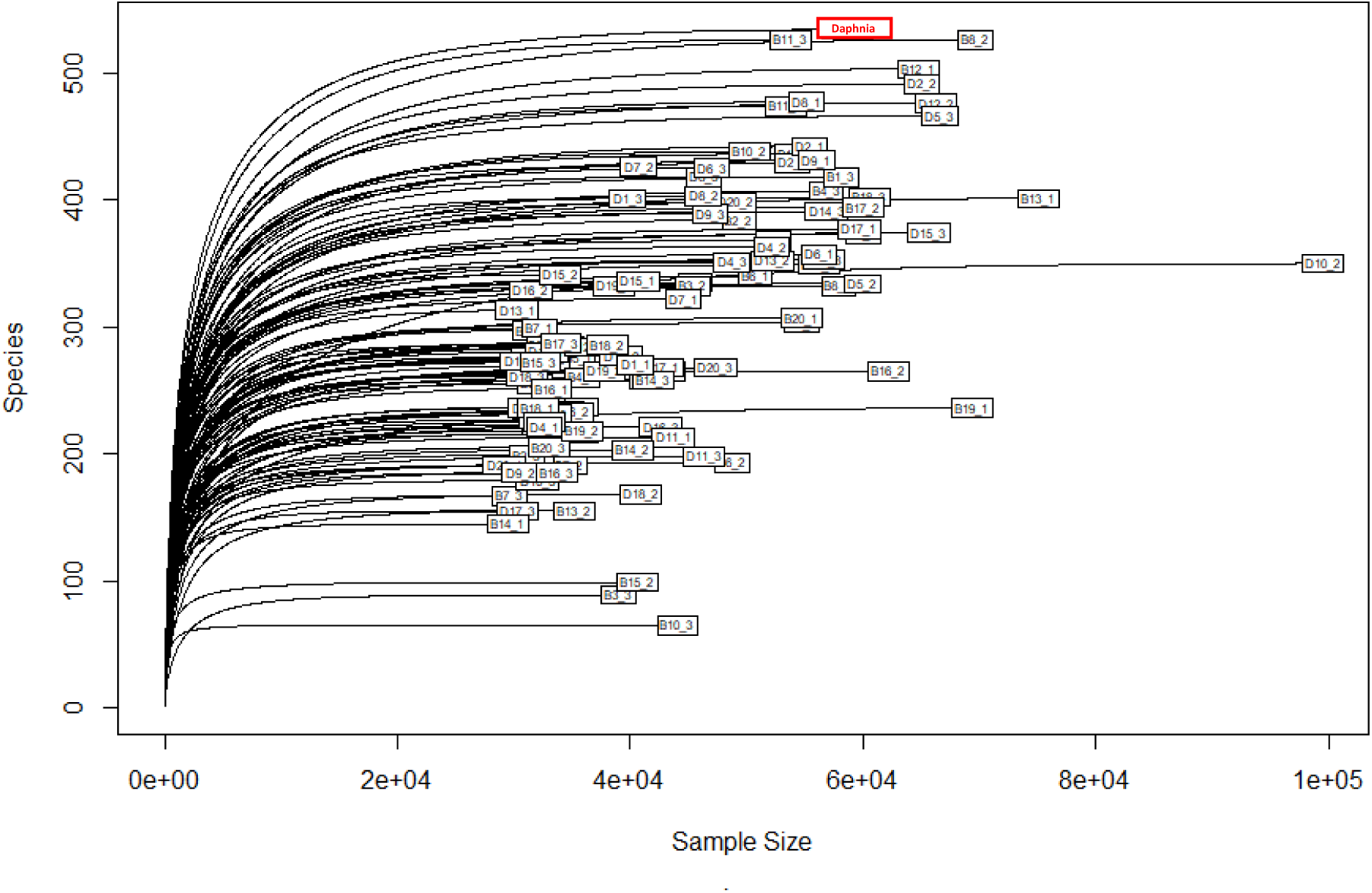
Rarefaction curves showing a good representation of the microbial communities in all samples. A composite Daphnia magna sample (10 individuals/sample) taken in the beginning of the exposure is shown in red.

**Table S5.** Predictors that were binarized for GLM and DESeq2 analyses.

| Predictor | Binary Threshold | Variable name |
| --- | --- | --- |
| **Experimental factors** |  |  |
| Suspended solids (SS in mg/l) | 0 vs 0.1-10 mg/l | SS (high/low) |
| MP (mass % of SS) | 0 vs pooled 1, 10 % | MP (yes/no) |
|  | 0, 1 vs 10% | MP <1 vs MP >1 |
| Agarose | 0 vs 1 |  |
| **Daphnia response** |  |  |
| Protein | Median protein content (0.011) | Protein (high/low) |
| Mortality (%) | <10% vs >10% | Mortality (high/low) |
| **PSD metrics** |  |  |
| D<sub>10</sub> |  |  |
| D<sub>50</sub> |  |  |
| D<sub>90</sub> |  |  |
| Kurtosis |  |  |
| Mode |  |  |
| Skewness |  |  |

**Figure S8.**
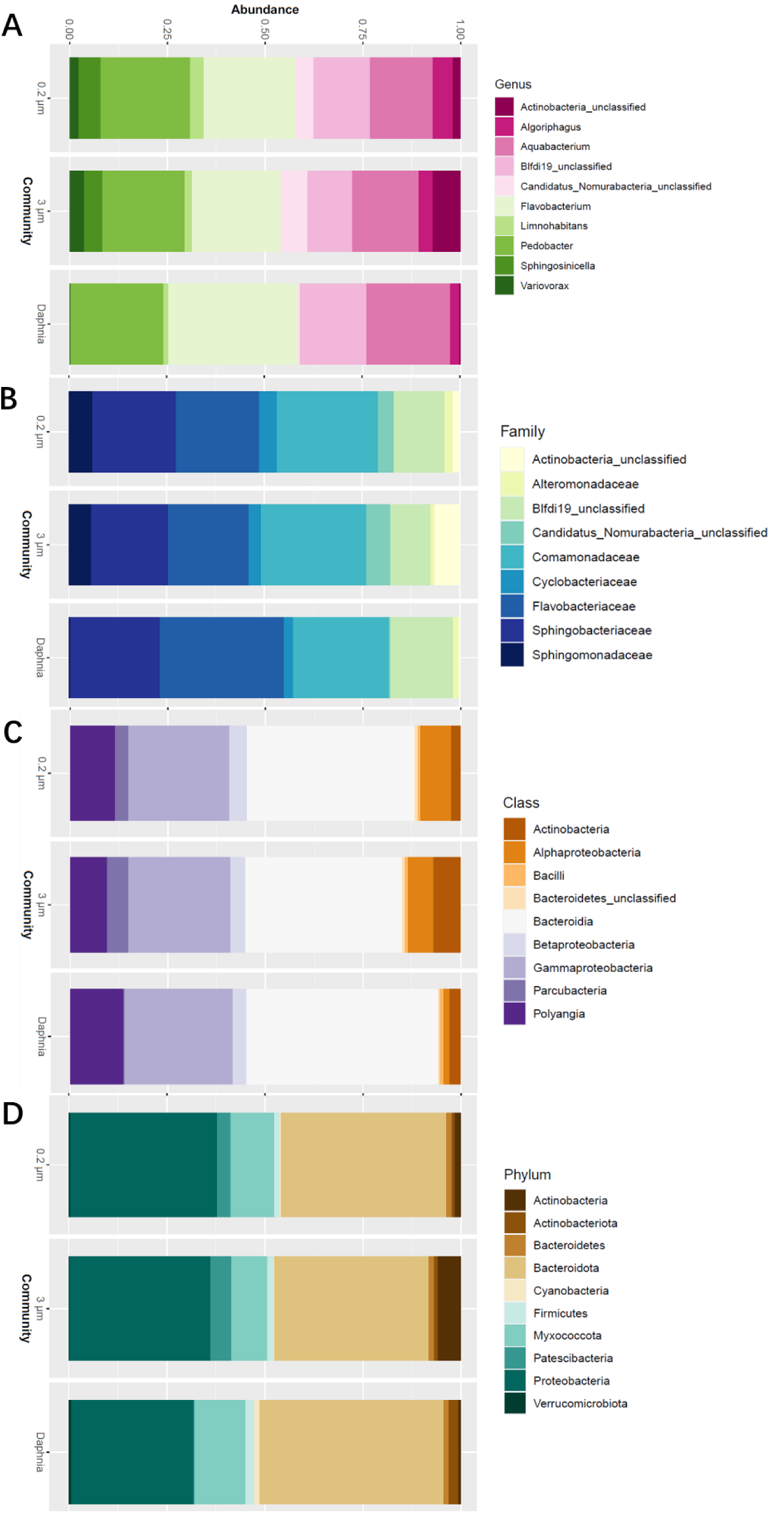
Overview of the relative abundances of bacterial taxa at four taxonomical levels (Phylum, Class, Family, Genus) in the samples of different size fractions and Daphnia magna

**Table S6.**
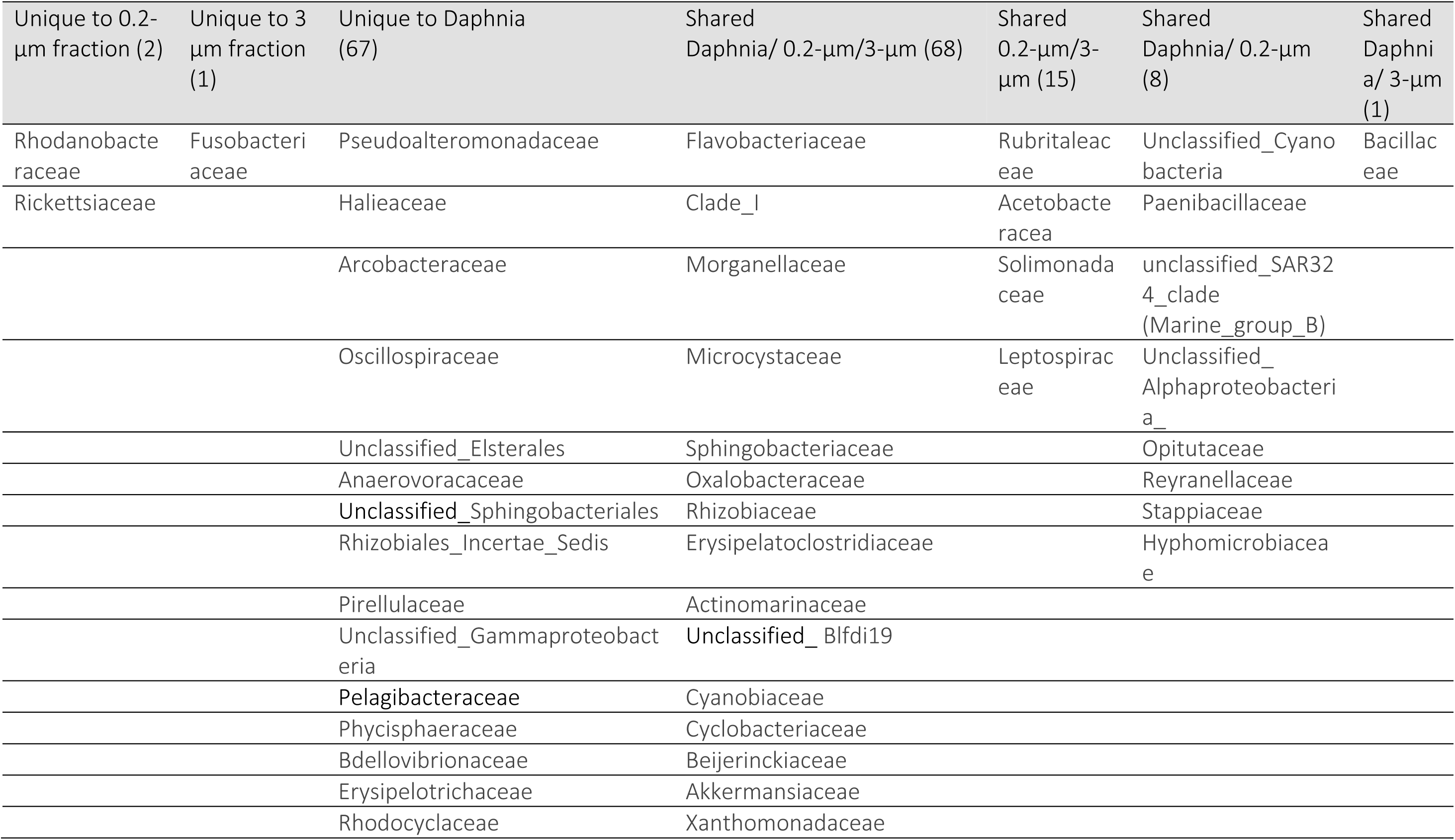
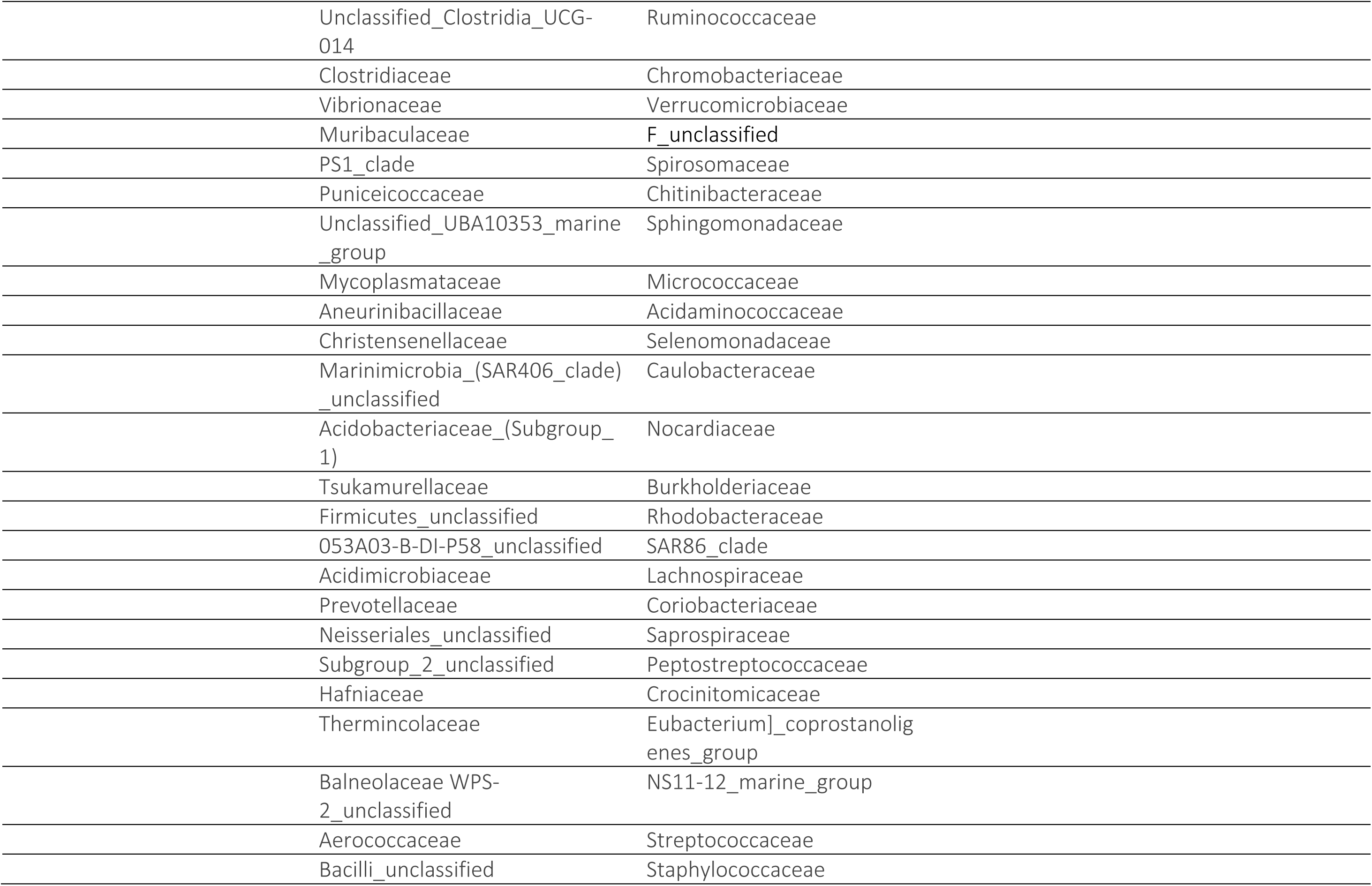
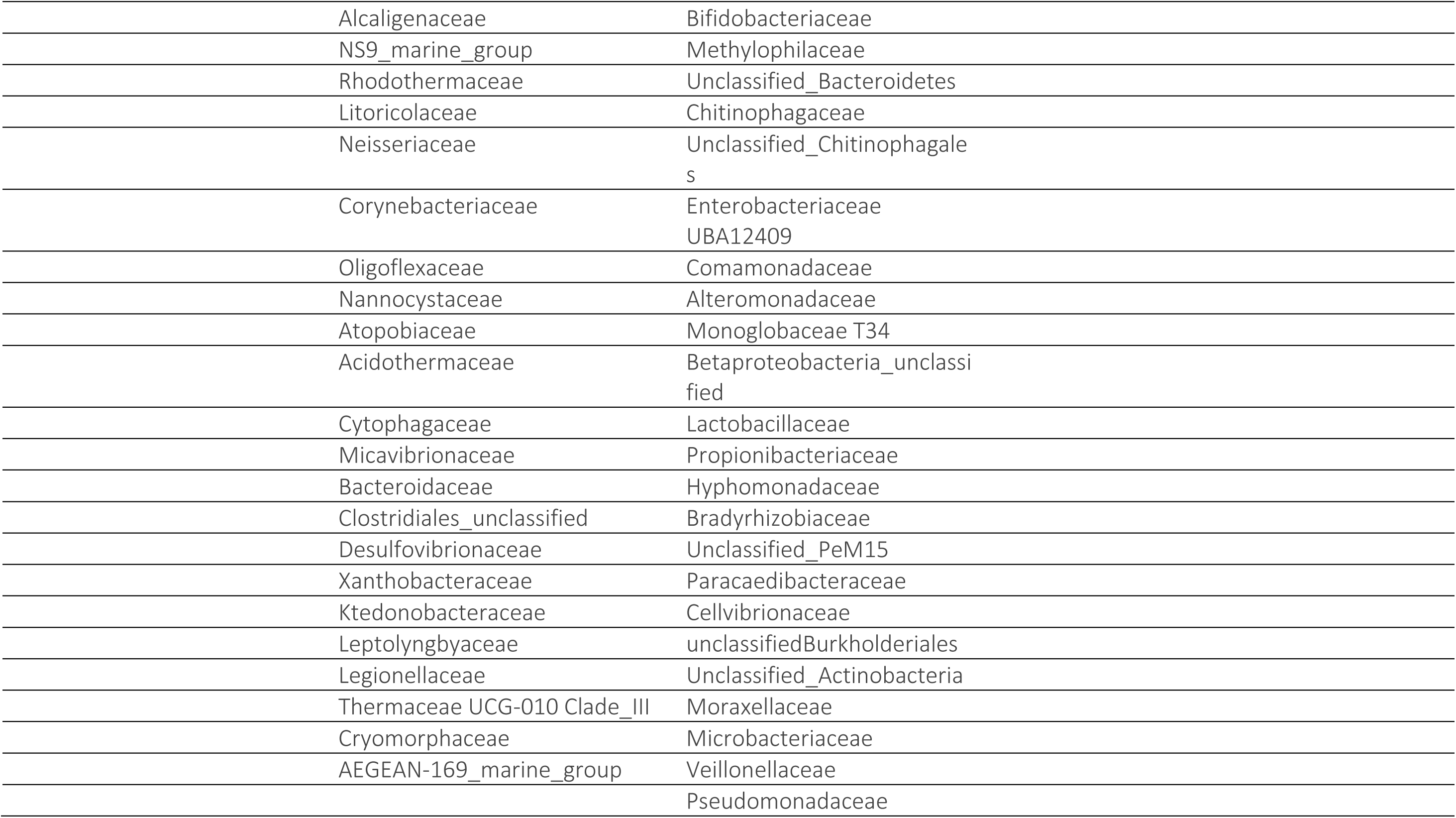
Taxa at the family level that were unique and those that were shared between the samples for the different size fractions (0.2-µm and 3-µm fractions representing non-adhering cells and biofilm, respectively) and Daphnia magna. See Figure 3 for the Venn diagram for visualization of these data.

**Table S7.** GLM output for the alpha diversity indices (Shannon and Fisher’s alpha) and experimental. The most parsimonious model was identified as a best-fit model with the fewest number of predictors using AIC.

|  | Estimate | Standard Error | Wald statistic | p-value |
| --- | --- | --- | --- | --- |
| Shannon |  |  |  |  |
| SS <1 | 0.0096 | 0.01603 | 0.36 | 0.5482 |
| Agarose | -0.0205 | 0.01529 | 1.8 | 0.1796 |
| Community | -0.0457 | 0.01534 | 8.870 | 0.0028 |
| MP (yes/no) | -0.0006 | 0.01639 | 0.002 | 0.9666 |

**Figure S9.**
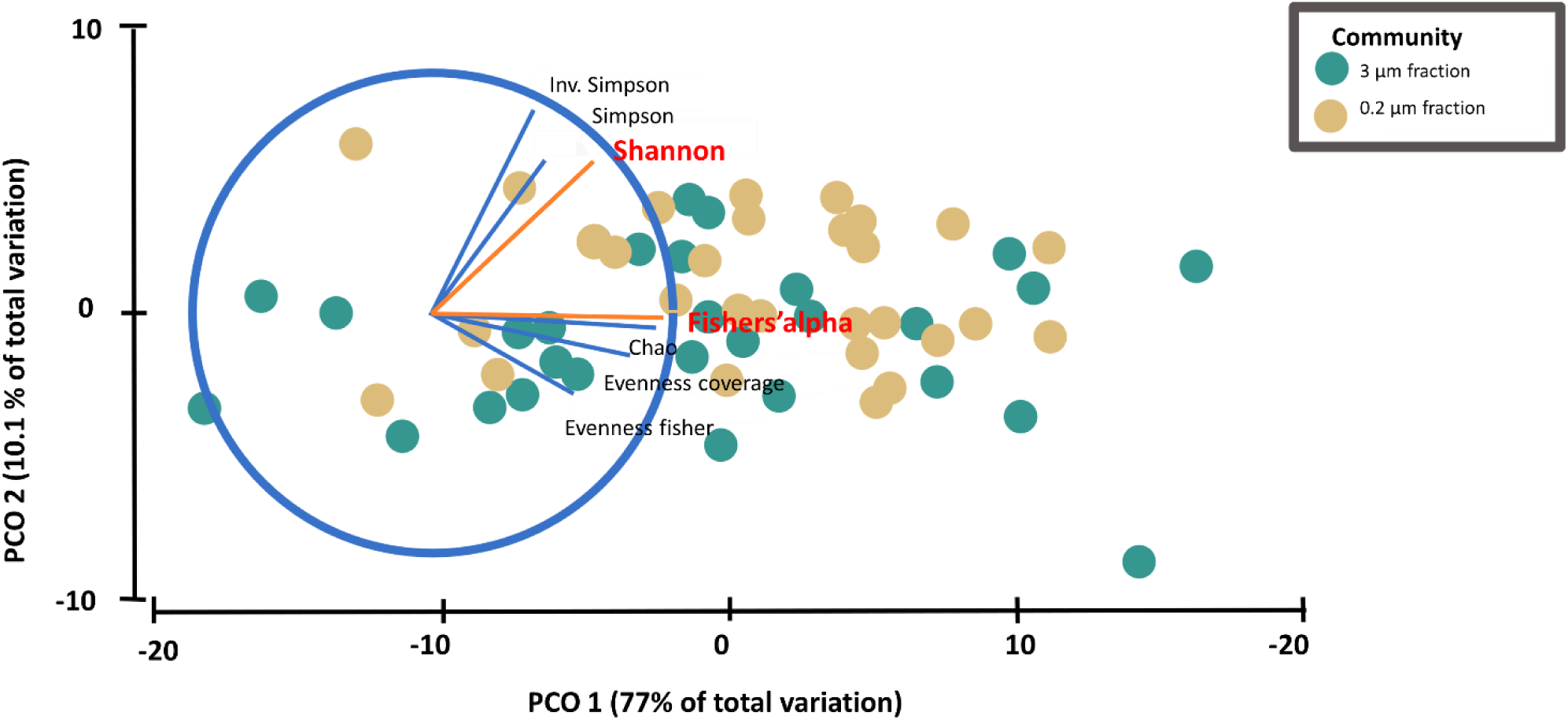
PCoA for various alpha diversity indices calculated for each sample of the 0.2- and 3-µm size fractions and based on the Bray-Curtis dissimilarity matrix. Fisher’s alpha and Shannon indices were selected as orthogonal and thus complementing each other for the alpha diversity analysis of the data in this study

**Figure S10.**
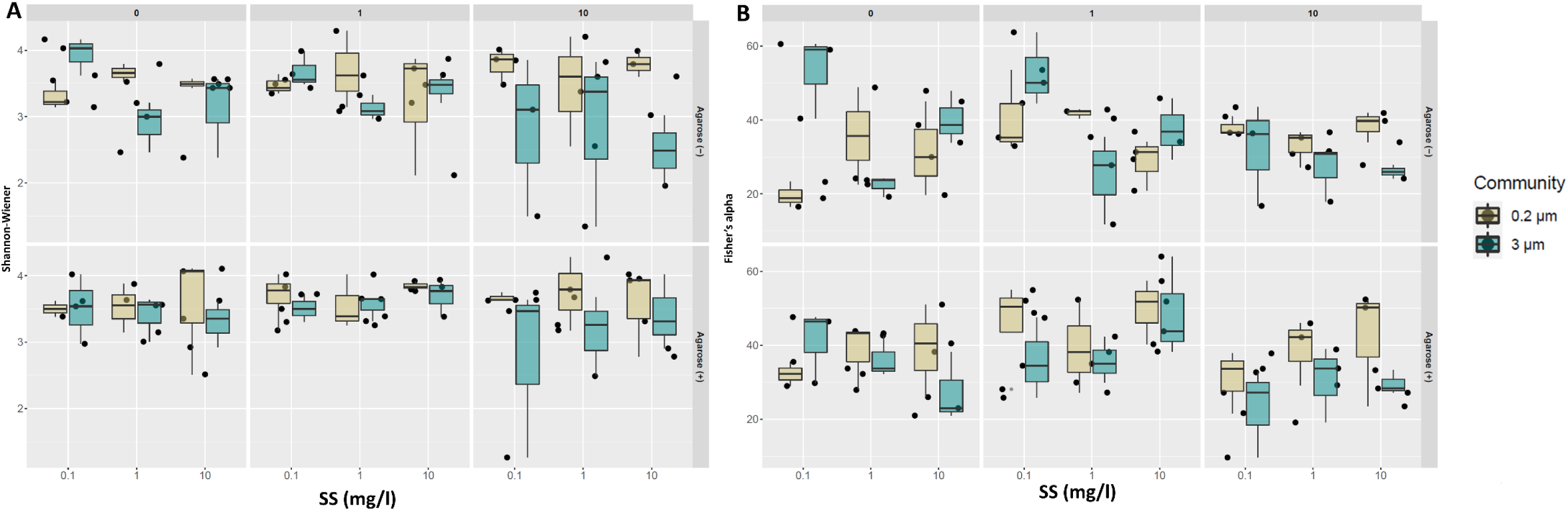
Boxplot showing Fisher.’s alpha (left) and Shannon diversity index (right) across different SS (x-axis) and MP% levels (grids), Agarose treatments are separated in rows (Agarose (-)/(+)).

**Table S8.**
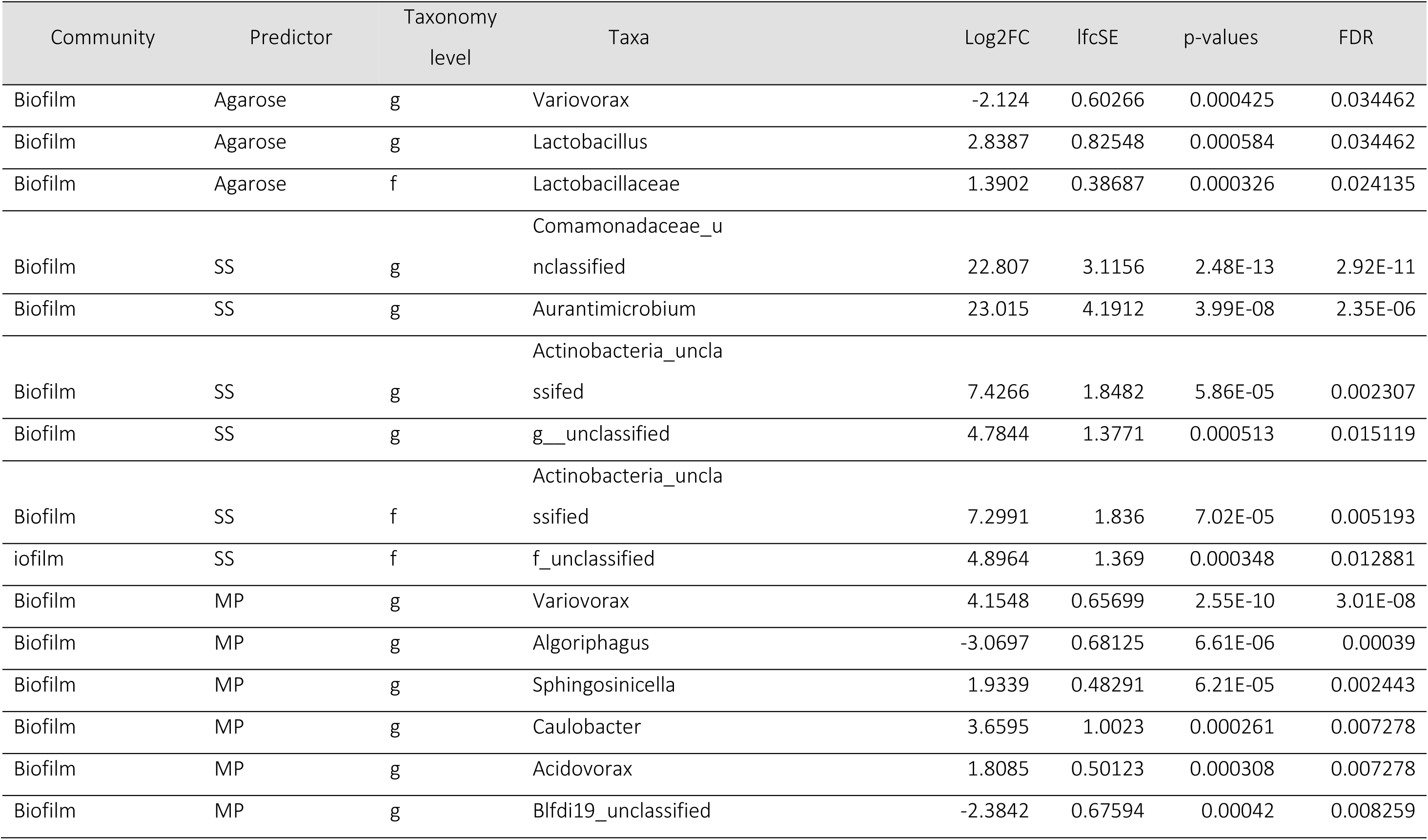
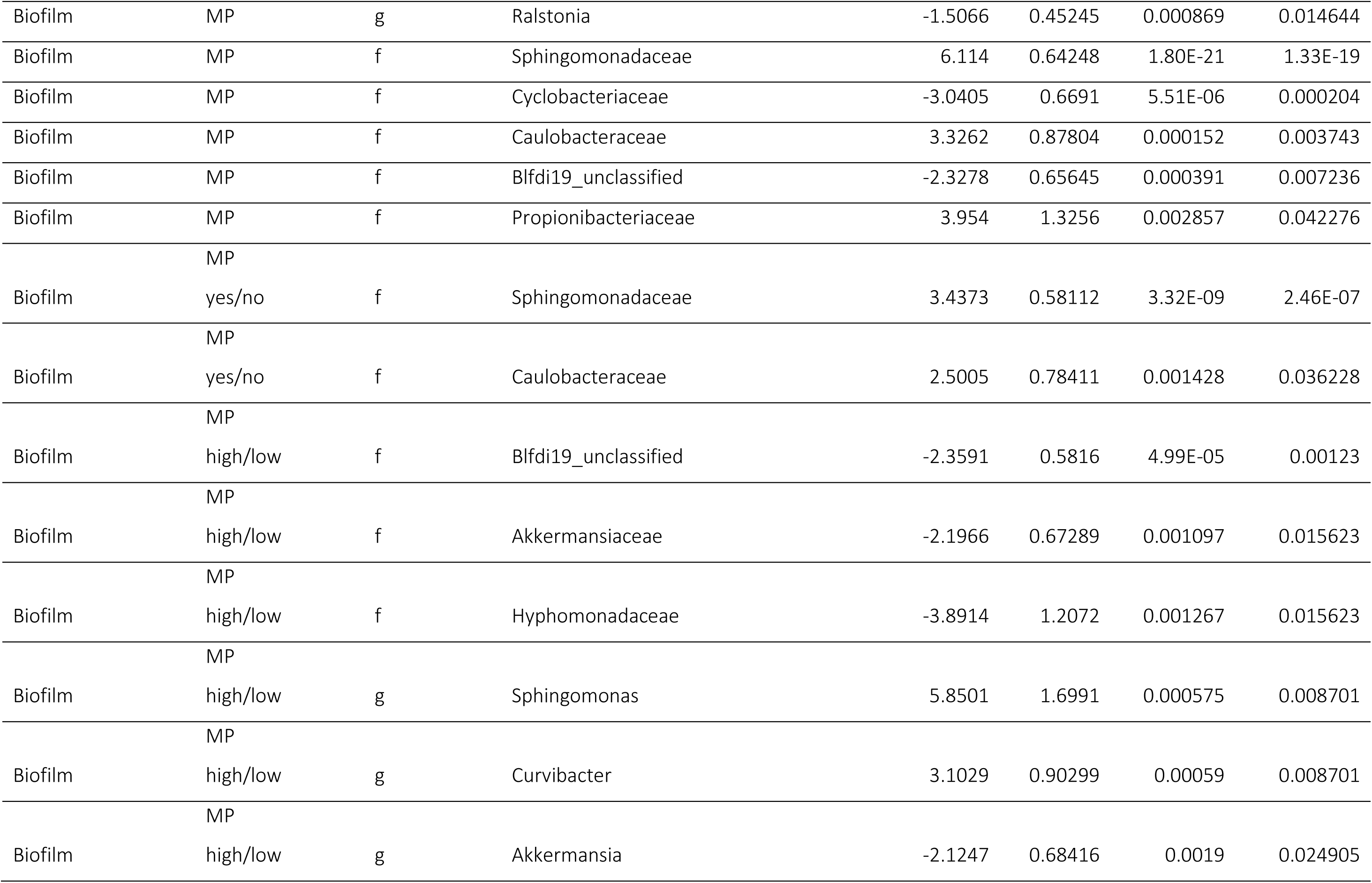
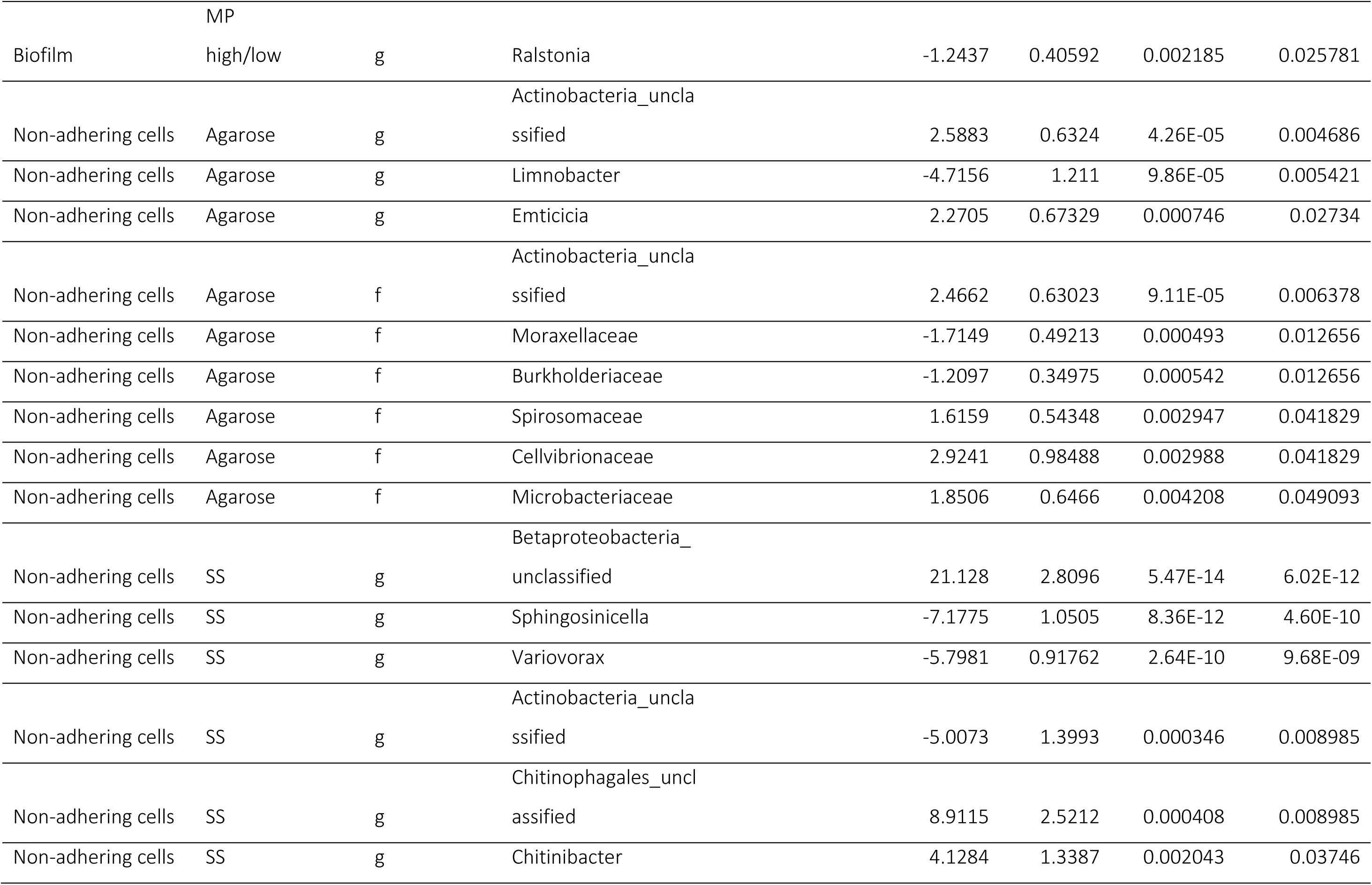
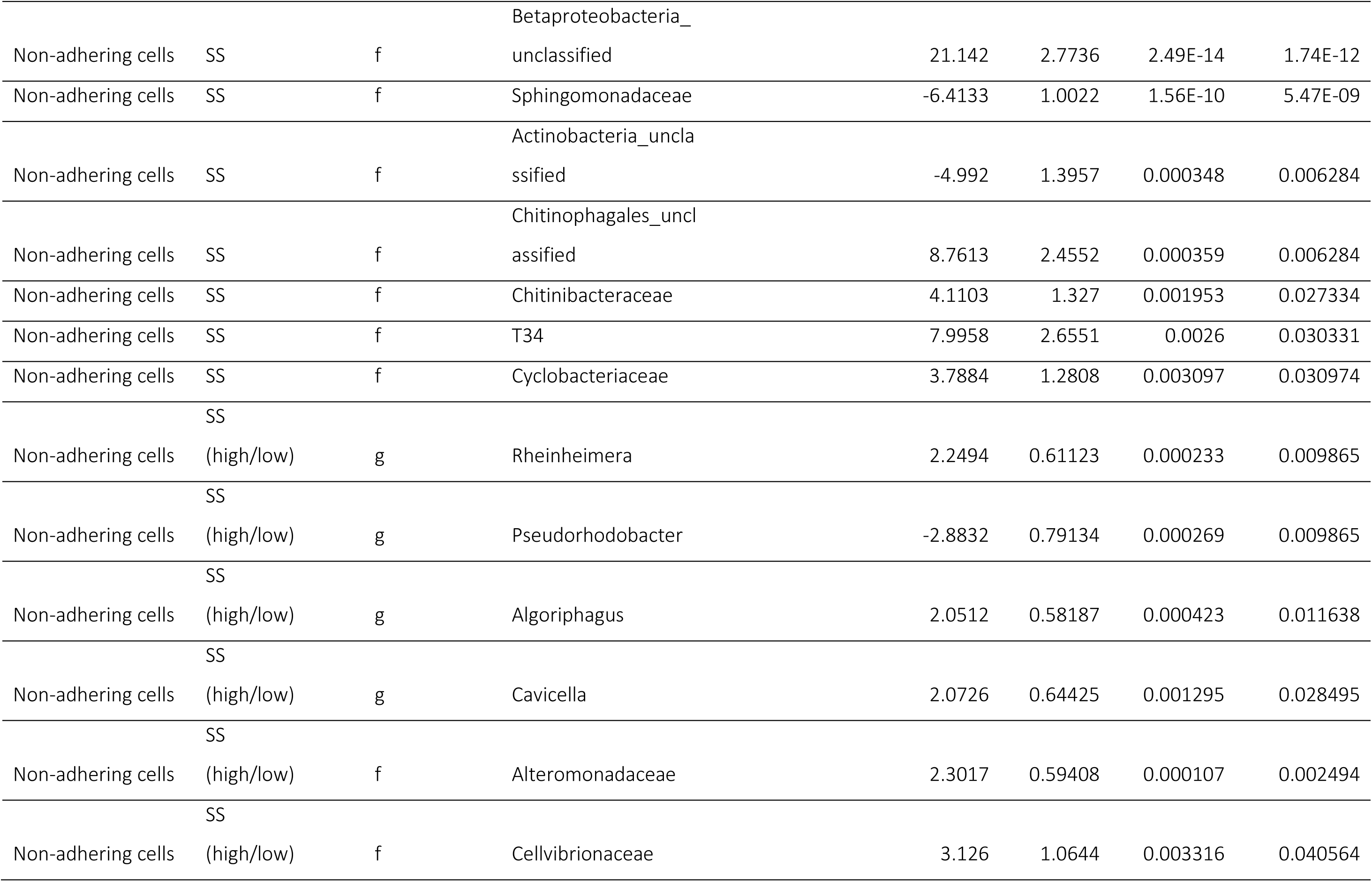
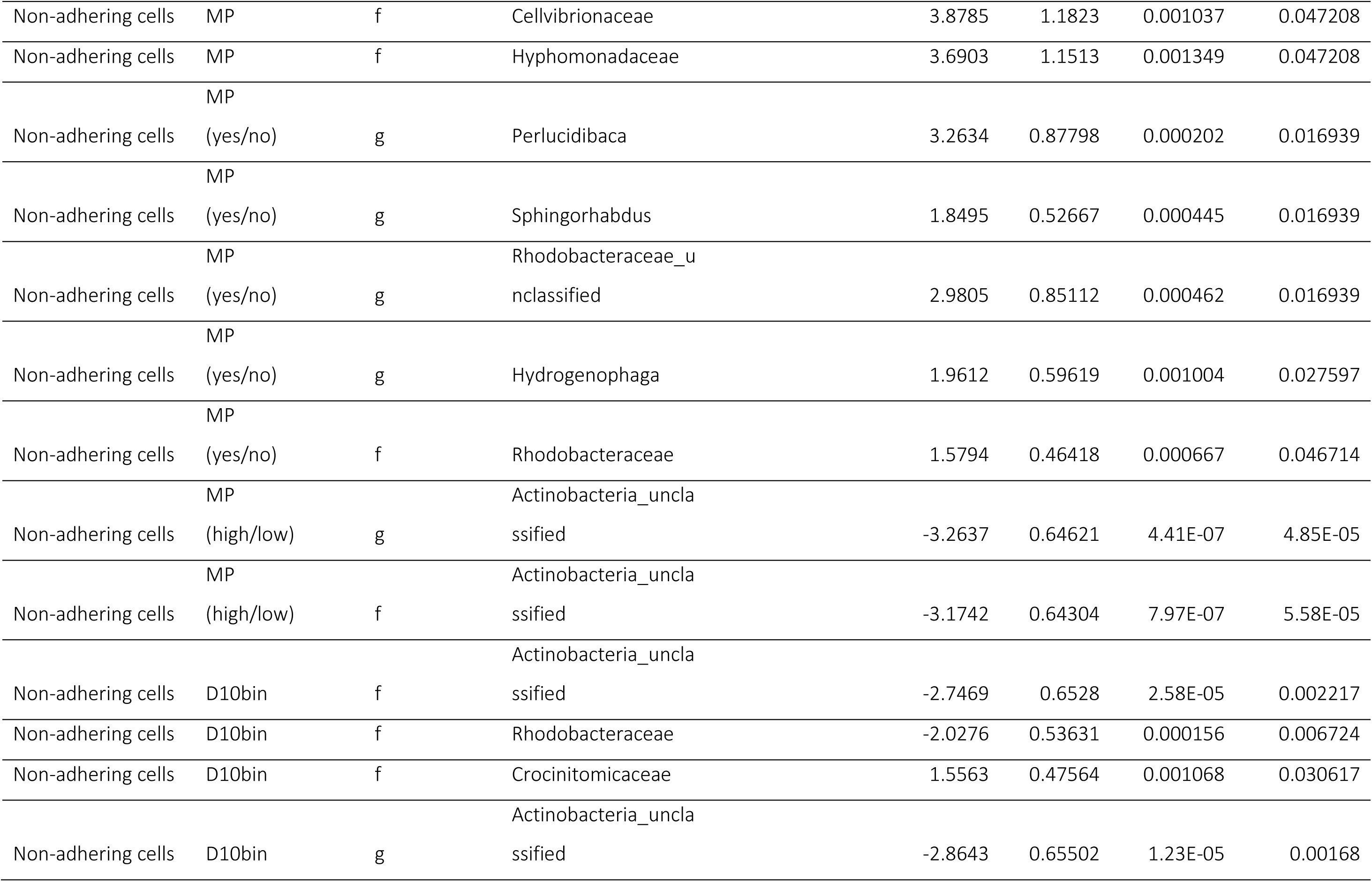
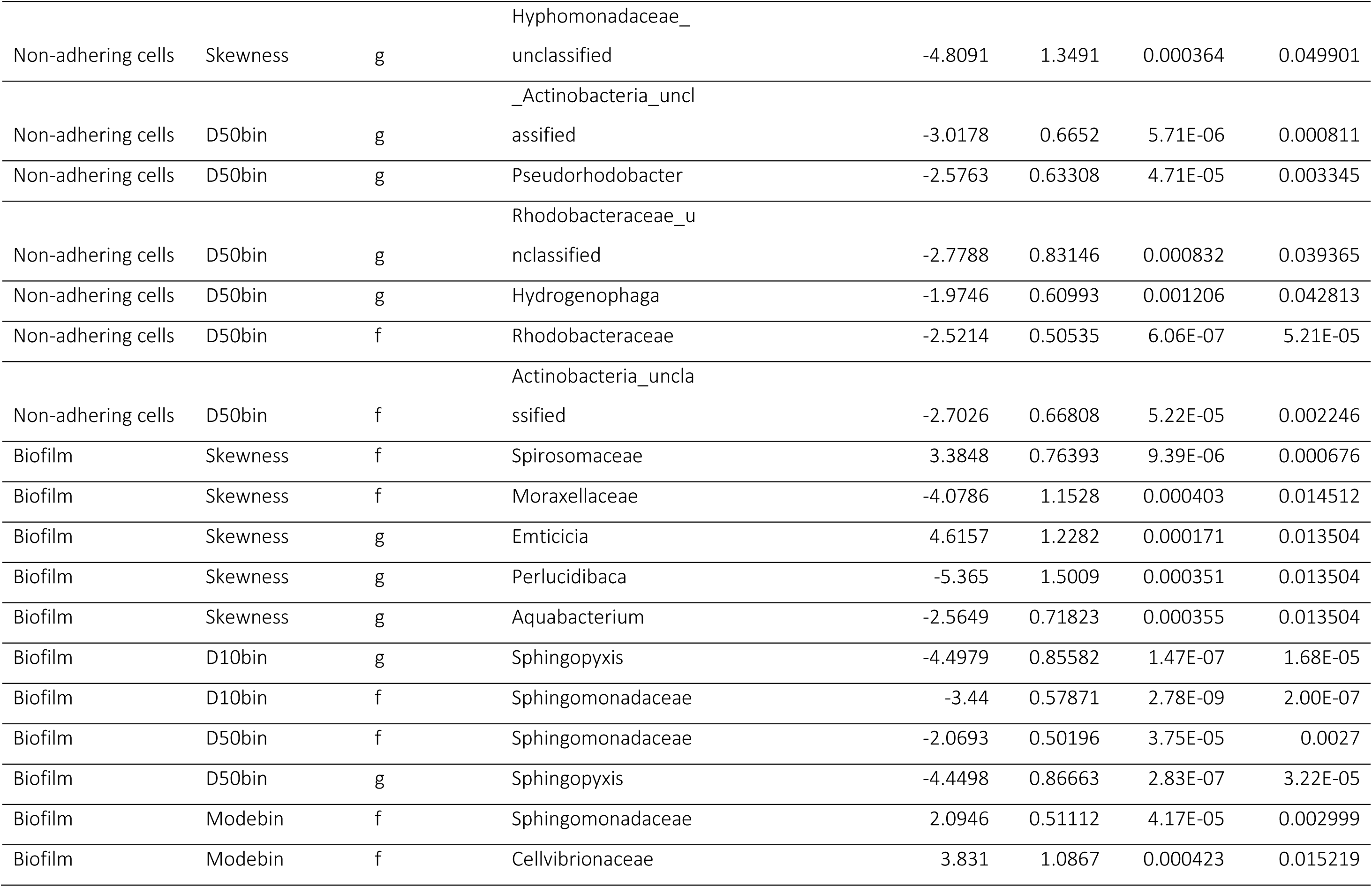
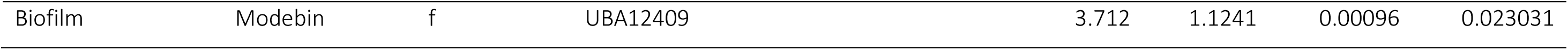
Differential abundance of taxa at family, genus and feature level for all experimental factors and PSD metrics identified by DESeq2 in biofilm (3-um fraction) and non-adhering cells (0.2-µm fraction). Significantly up- and downregulated taxa across communities, experimental factors and PSD metrics are shown.

**Figure S11.**
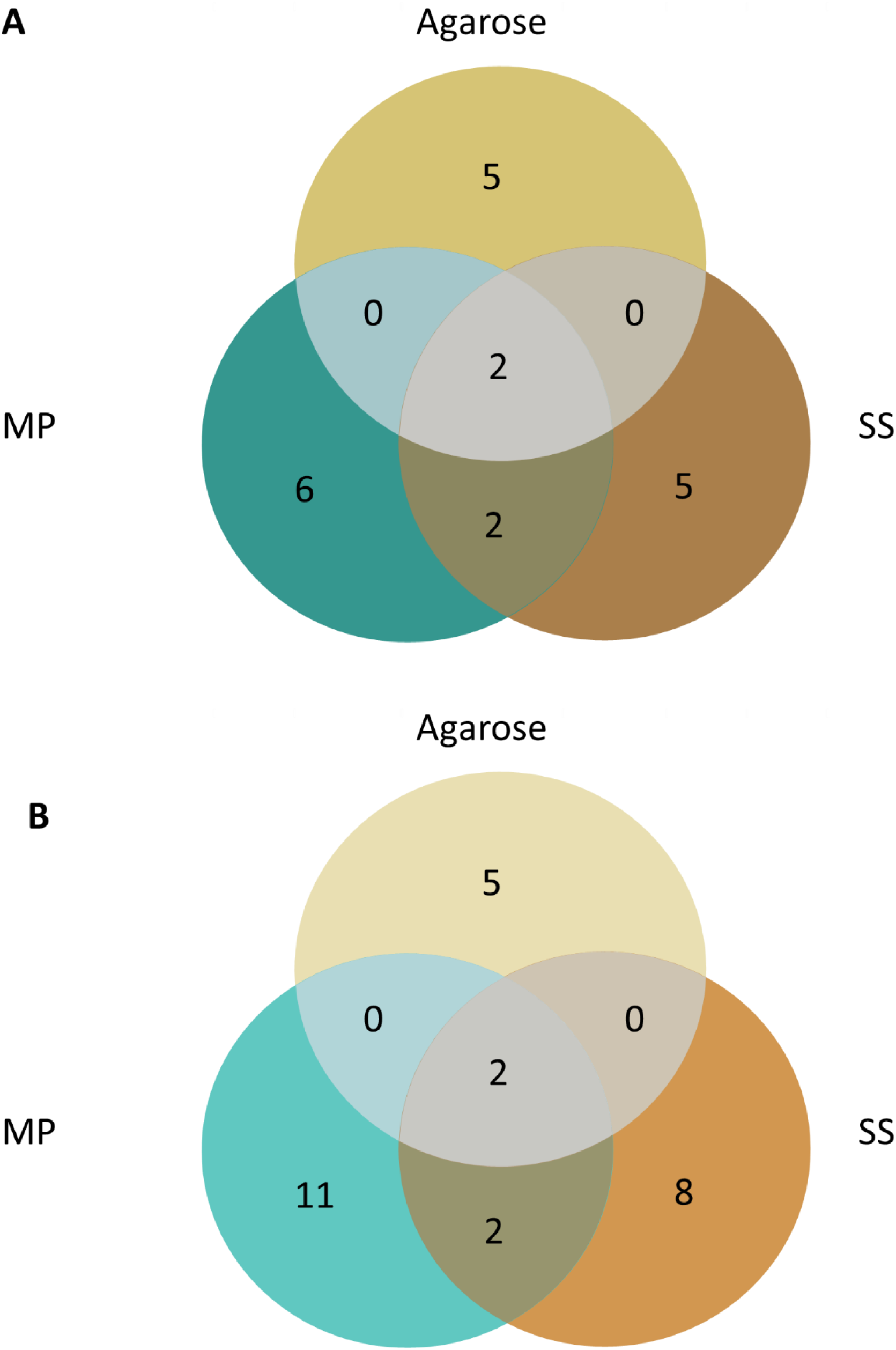
Venn diagram for shared and unique families (A) and genera (B) affected by the experimental factors and identified by DESeq2. See table X for the summary of the data.

**Table S9.**
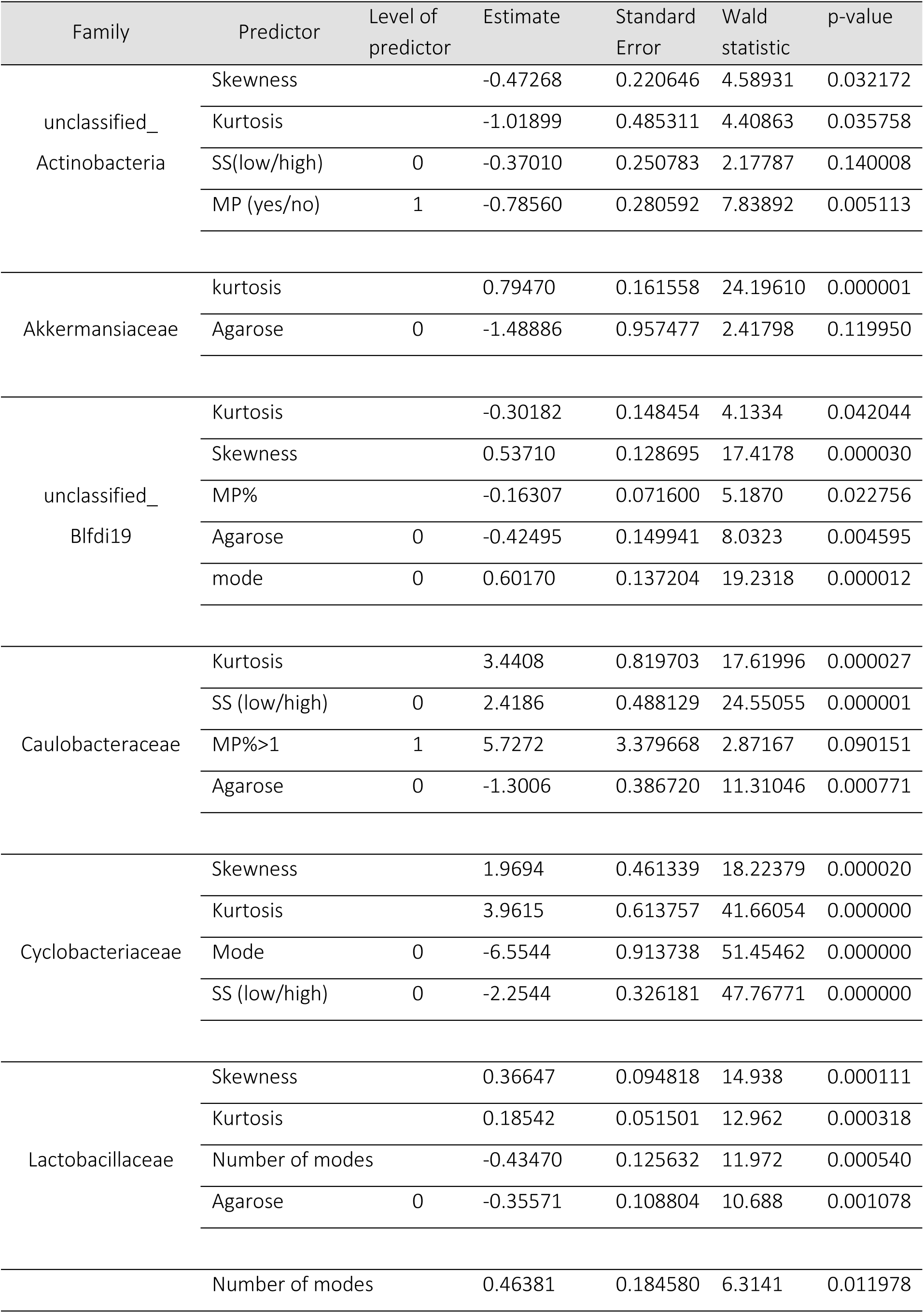
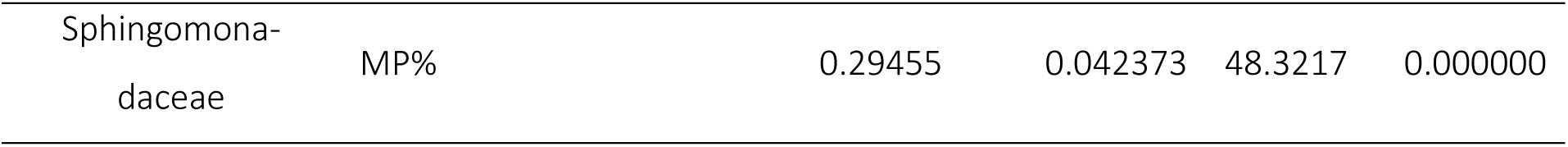
Best-fit GLM for the relative abundance of the affected families as a function of the experimental factors and PSD metrics.

## Notes

### Competing Interest Statement

The authors have declared no competing interest.

### Funding

- VINNOVA
- Bonus

